# Impaired dynamics of prefrontal beta-band networks during cognitive action control in Parkinson’s disease

**DOI:** 10.1101/2021.12.12.472263

**Authors:** Joan Duprez, Judie Tabbal, Mahmoud Hassan, Julien Modolo, Aya Kabbara, Ahmad Mheich, Sophie Drapier, Marc Vérin, Paul Sauleau, Fabrice Wendling, Pascal Benquet, Jean-François Houvenaghel

**Affiliations:** Univ Rennes, LTSI - U1099, F-35000 Rennes, France; Azm Center for Research in Biotechnology and Its Applications, EDST, Lebanese University, Beirut, Lebanon; MINDig, F-35000 Rennes, France; School of Engineering, Reykjavik University, Iceland; Neurokyma, F-35000 Rennes, France; CIC INSERM 1414, Rennes, France; Neurology Department, Pontchaillou Hospital, Rennes University Hospital, France; ‘Behavioral and Basal Ganglia’ Research Unit, University of Rennes 1-Rennes University Hospital, France; Neurophysiology department, Rennes University Hospital, France

**Keywords:** Functional connectivity, Networks, Dynamics, high density EEG, Simon task, Parkinson’s disease

## Abstract

Among the cognitive symptoms that are associated with Parkinson’s disease (PD), alterations in cognitive action control (CAC) are commonly reported in patients. CAC enables the suppression of an automatic action, in favor of a goal-directed one. The implementation of CAC is time-resolved and arguably associated with dynamic changes in functional brain networks. However, the electrophysiological functional networks involved, their dynamic changes, and how these changes are affected by PD, still remain unknown. In this study, to address this gap of knowledge, 21 PD patients and 10 healthy controls (HC) underwent a Simon task while high-density electroencephalography (HD-EEG) was recorded. Source-level dynamic connectivity matrices were estimated using the phase-locking value in the beta (12-25 Hz) and gamma (30-45 Hz) frequency bands. Temporal independent component analyses were used as a dimension reduction tool to isolate the group-specific brain network states that were dominant during the task. Typical microstate metrics were quantified to investigate the presence of these states at the subject-level. Our results first confirmed that PD patients experienced difficulties in inhibiting automatic responses during the task. At the group-level, HC displayed a significant functional network state that involved typical CAC-related prefrontal and cingulate nodes (e.g., inferior frontal cortex). Both group- and subject-level analyses showed that this network was less present in PD to the benefit of other networks involving lateralized temporal and insular components. The presence of this prefrontal network was associated with decreased reaction time. In the gamma band, two networks (fronto-cingulate and fronto-temporal) followed one another in HC, while 3 partially overlapping networks that included fronto-temporal, fronto-occipital and cross-hemispheric temporal connections were found in PD. At the subject-level, differences between PD and HC were less marked. Altogether, this study showed that the functional brain networks observed during CAC and their temporal changes were different in PD patients as compared to HC, and that these differences partially relate to behavioral changes. This study also highlights that task-based dynamic functional connectivity is a promising approach in understanding the cognitive dysfunctions observed in PD and beyond.

**Highlights:**

- Cognitive action control is associated with dynamic functional networks
- Prefrontal and cingulate beta connectivity are prominent in healthy controls
- PD patients have different dynamic networks in which prefrontal nodes are absent
- The occurrence of prefrontal beta networks was associated with a decreased reaction time
- Functional networks in the gamma band were temporally organized in HC, but overlapping in PD patients

## 1. Introduction

Parkinson’s disease (PD) is associated with a broad spectrum of symptoms. Although it is mostly known for its deleterious effects on motor function, such as bradykinesia, rigidity and resting tremor (Hayes, 2019); cognitive impairments associated with PD also have a significant impact on quality of life (Lawson et al., 2016). One of the major cognitive difficulties found in PD patients is the impairment in efficient and fast adaptation to environmental changes. More specifically, PD patients typically show alterations in cognitive action control (CAC), a sub-process of cognitive control that enables the suppression of automatic responses in favor of goal-directed voluntary actions (as considered in light of the dual-route and activation-suppression models, see Hommel and Wiers, 2017). PD patients consistently show alterations regarding this specific function compared to healthy controls (HC), although the nature of the impairment is somewhat inconsistent between studies (Cagigas et al., 2007; Duprez et al., 2017; Falkenstein et al., 2006; Wylie et al., 2010, 2005).

Importantly, even at the prodromal stages of the disease, accumulation of *α*-Synuclein results in synaptic and axonal dysfunctions (Bridi and Hirth, 2018). This could explain the presence of cognitive impairment that is sometimes already present at the time of diagnosis. Indeed, integrity and efficiency of synaptic communication is arguably important for processes requiring fast decision making such as CAC. Several studies have investigated the correlations between brain activity on the one hand, and cognitive changes on the other hand, using a variety of neuroimaging modalities (fMRI, PET, (M)/EEG). For instance, it is now undisputed that conflict resolution is associated with activity in the dorso-lateral prefrontal cortex (DLPFC), the inferior frontal cortex (IFC), the pre-supplementary motor area (pre-SMA), the anterior cingulate cortex, and the subthalamic nucleus (Ridderinkhof et al., 2011). In addition to this localizationist approach, studying the interaction between brain regions would help in specifying how neurodegenerative diseases such as PD alter cognitive functions. Indeed, a large body of evidence now shows that cognitive functioning emerges from the communication of distant brain regions (Bassett and Sporns, 2017) and that alterations in these brain networks have been associated with neurological disorders (Fornito et al., 2015). Brain networks can be studied, among available techniques, via the estimation of functional connectivity (FC) between brain areas, as estimated from electrophysiological ((M)EEG) or metabolic (MRI, PET) signals. FC is not in itself a direct measure of communication between brain areas, but rather reflects statistical dependencies of brain activity between different regions. (M)EEG FC is particularly interesting since it is usually inferred through the phase synchronization of neural oscillations, a mechanism that has been proposed to facilitate communication between neuronal assemblies (Fries, 2015).

So far, most studies have evaluated FC in PD using resting-state fMRI (Baggio et al., 2015, 2014; Lopes et al., 2017; Skidmore et al., 2011; Wolters et al., 2019). The literature focusing on FC evaluated by (M)EEG and/or during cognitive tasks in PD is scarcer, although it would surely provide valuable insights on the neurophysiological alterations caused by the disease and its relationship with cognitive symptoms. While (M)EEG does not benefit from the same spatial resolution as fMRI, recent advances in cortical source reconstruction now enable the inference of cortical area long-range FC (Hassan and Wendling, 2018). One major advantage of (M)EEG techniques is their excellent temporal resolution (millisecond scale), which is fundamental when studying cognitive processes that are inherently fast and dynamic. For instance, CAC is a process that has obvious dynamic properties at the behavioral level: action selection and suppression are time-resolved processes allowing conflict resolution, and these aspects were usually masked by focusing on task-averaged behavioral performances in princeps studies (van den Wildenberg et al., 2010).

Similarly to cognitive processes, FC is also time-dependent and brain networks dynamically rearrange themselves during resting state and tasks (Baker et al., 2014; Bola and Sabel, 2015; de Pasquale et al., 2016; Hassan et al., 2015; Kabbara et al., 2021; O’Neill et al., 2018, 2017) with consequences on behavior (Allen et al., 2018). Several new methods now allow investigating how inter-regional communication varies with time (Sizemore and Bassett, 2018; Tabbal et al., 2021). Such methods would tremendously help in understanding how dynamic cognitive processes such as CAC unravel, and how those processes are impaired in neurodegenerative diseases such as PD. In this study, we hypothesize that PD is associated with changes in the dynamic properties of brain networks during CAC. We use the example of a classic conflict task, the Simon task, paired with high-density EEG (HD-EEG, 256 channels) reconstructed at the cortical source level in a group of PD patients and in HC. We combine the calculation of dynamic FC matrices with a dimension reduction method (independent component analysis, ICA) and micro-state metrics to investigate the time-varying changes in brain networks at the group- and subject-level. This allowed us to identify and characterize the frequency-dependent networks underlying the difference in brain network dynamics between PD patients and HC.

## 2. Materials and methods

### 2.1. Participants

Twenty-one patients (10 males, 11 females) diagnosed with idiopathic PD (Hughes et al., 1992), aged between 48-69 years (mean=59.4, std=6.7) and 10 HC (4 males and 6 females), aged between 45-57 years (mean=52.8, std=4.3) participated in this study. Patients with major cognitive deficits (Montreal Cognitive Assessment [MoCA], <22) or with other past or present neurological (other than PD) or psychiatric pathology, as well as patients with deep brain stimulation, were not included in the study. The same criteria were used for HC. Global cognitive functioning was assessed in all participants using the MoCA (Nasreddine et al., 2005). Additional standardized tests were assessed for PD patients, representing several cognitive abilities. These tests included: the Symbol Digit Modalities Test (SDMT; Smith, 1982), the Digit span test (Wechsler, 1981), the Stroop test (Stroop, 1935), the judgment of line orientation test (Benton et al., 1978), the Boston naming test (Graves et al., 2004), as well as semantic fluency (animal names generation task in 60 seconds), and phonemic fluency (words generation task in 60 seconds). All HC were recruited from the general population during public conferences and through participation calls. Patients were recruited at the University Hospital of Rennes, France during hospitalization for their usual care. All participants provided informed consent for participation in the study, which was approved by a national ethics committee review board (CPP ID-RCB: 2019-A00608-49; approval number: 19.03.08.63626).

### 2.2 Experimental Task

In this study, we used a color version of the Simon task (Simon and Rudell, 1967) to investigate the changes in CAC associated with PD. Participants were seated at a distance of 80 cm from a 22 inches’ computer screen. At the beginning of a trial, a central dark fixation cross was presented on a white screen, during a variable period (pseudo-randomly defined from 1750 ms to 2170 ms). Then, a blue or a yellow circle (3.9 cm diameter) was displayed either on the right or left side during 200 ms (Figure 1). Participants were asked to press a left or right button according to the color of the circle displayed, while ignoring its location (color/side mapping was counterbalanced across participants). Participants had to respond within 1000 ms after stimulus offset. The location side of the colored circle stimulus could match (congruent) or not (incongruent) the side of the correct button press associated with the color. For example, when the color “blue” is mapped to the “right” button, the trial is congruent when a blue circle appears on the right side, and incongruent when it appears on the left side (Figure 1). After a 60-trials training session, participants performed 10 blocks of 60 trials, with short pauses between blocks and a longer pause every three blocks to check EEG electrodes’ impedance. In total, 600 trials were performed with 300 congruent and 300 incongruent trials with a pseudo-randomized display.

**Figure 1.**
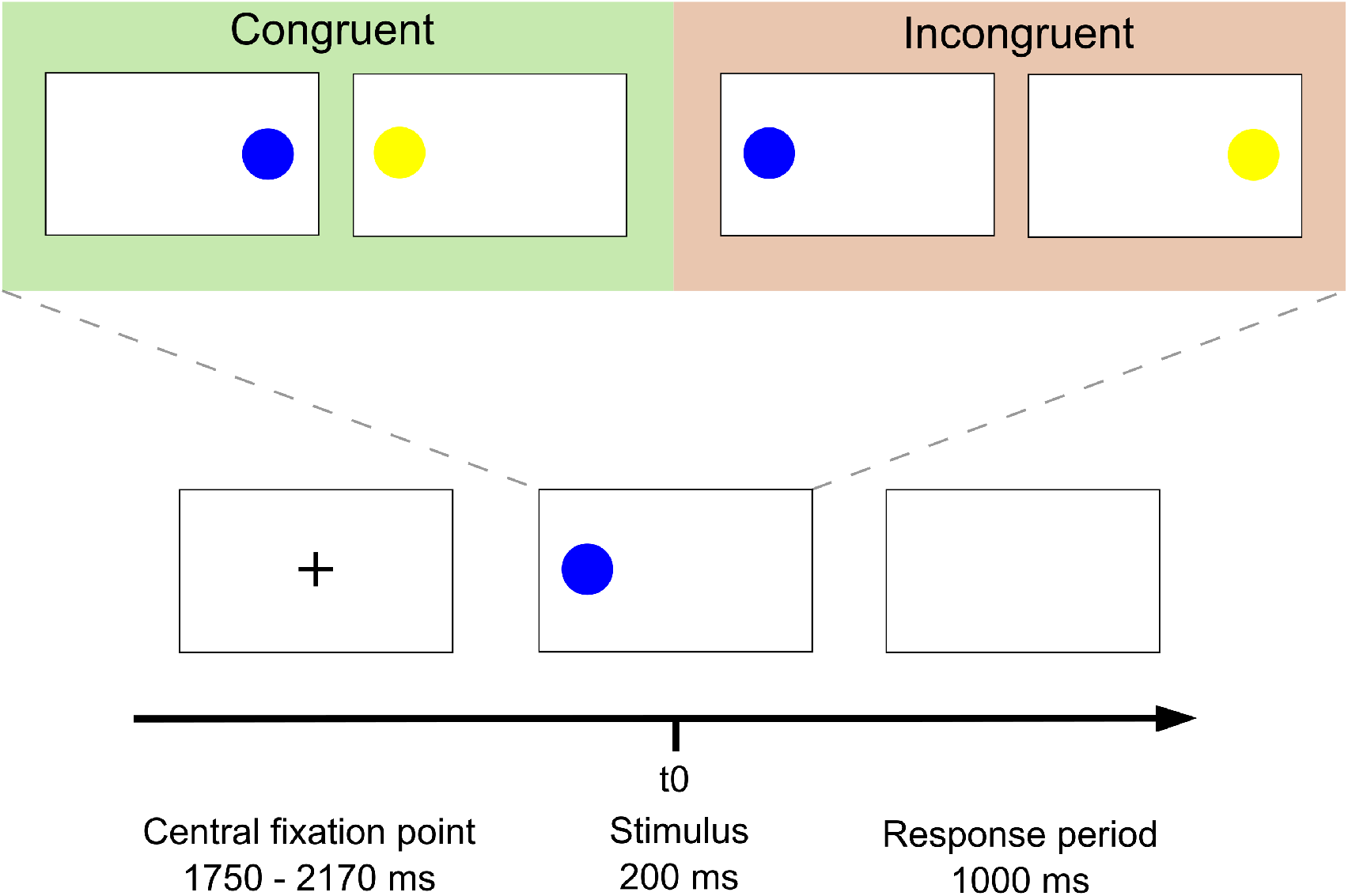
Overview of the Simon Task. The central fixation point was displayed randomly from 1750 to 2170 ms, with a 30 ms step. Then, the stimulus display lasted 200 ms. Participants had 1000 ms to answer by pressing the button. Two conditions could occur: congruent, when color and location of the circle led to the same response; and incongruent, when color and location didn’t lead to the same response.

### 2.3 Data acquisition and Preprocessing

EEG signals were recorded using a HD-EEG system (EGI, Electrical Geodesic Inc., 256 channels), with a sampling frequency of 1000 Hz. 199 electrodes were kept, removing most of the jaw and neck electrodes, as shown in the channel file available in the GitHub repository (see section 2.8). EEG preprocessing was performed manually using the Brainstorm toolbox (Tadel et al., 2011). Preprocessing was performed as follows. First, DC offset removal was applied. Second, a notch filter at 50 Hz and a band-pass filter of 1-100 Hz were applied. Third, signals were visually inspected, and bad channels were removed before being interpolated using Brainstorm’s default parameters. Fourth, Independent Component Analysis (ICA) was used to remove eye blinks and muscle artifacts. Fifth, we segmented the recorded signals into epochs from -700 ms to 1200 ms relative to the stimulus onset. Finally, a visual inspection was performed to manually reject epochs with excessive remaining noise. For EEG analyses, we focused on correct incongruent trials only, since they are associated with efficient control in a condition of strong conflict. As a result, out of originally 300 incongruent trials, 174 incongruent trials per subject on average (STD=49.2) were kept for further analyses because of the removal of errors, the absence of response, or because of remaining artifacts. After EEG preprocessing, several steps were applied to identify the dynamic brain network states in each group (HC and PD), as summarized in Figure 2 and explained below.

**Figure 2.**
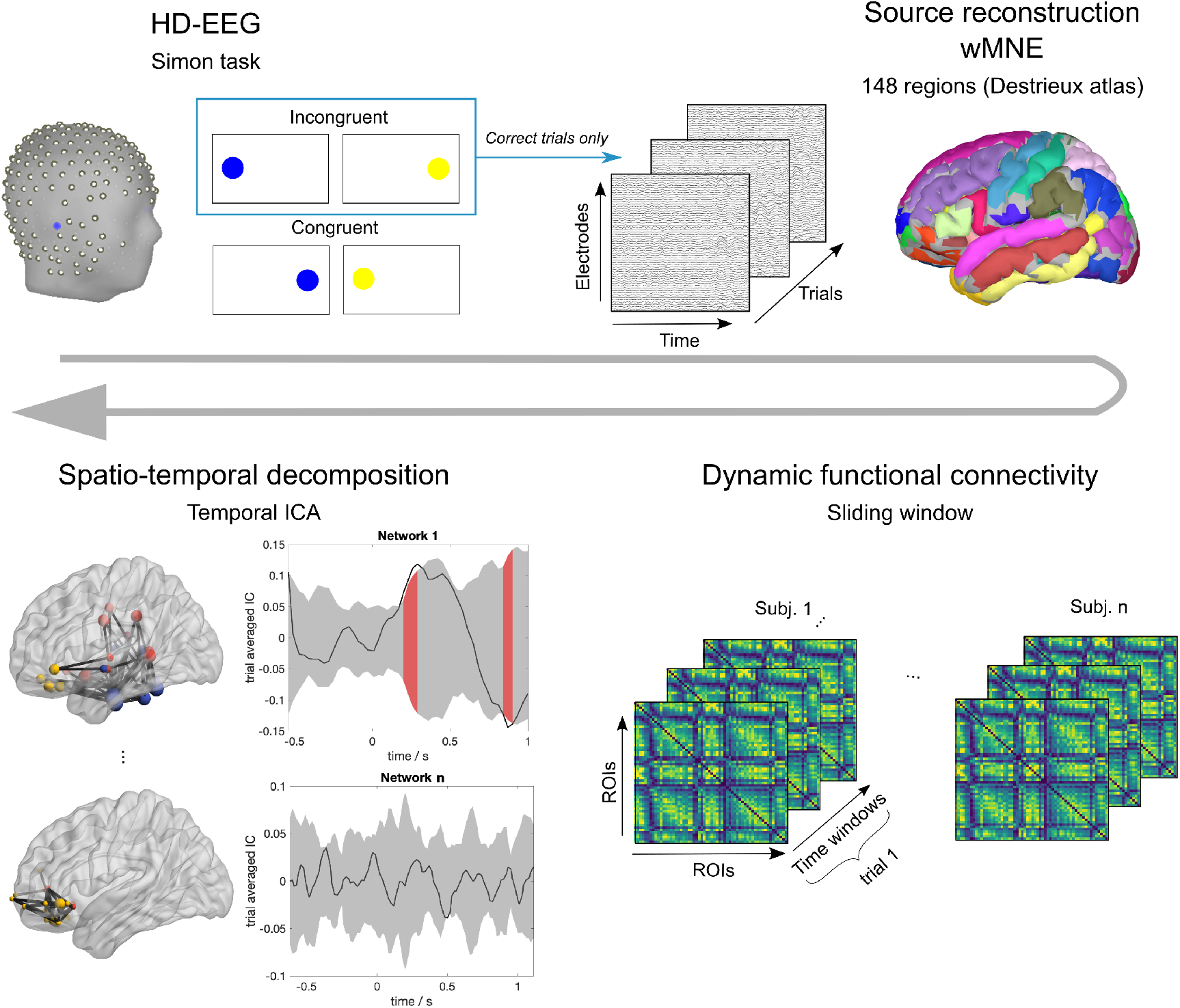
Dynamic FC pipeline used in our study. First, HD-EEG data was recorded during the Simon task (only correct incongruent trials were considered). After preprocessing, cortical-level sources were reconstructed using the weighted Minimum Norm Estimate (wMNE) and the Destrieux atlas (148 Regions of Interests, ROIs). Then, dynamic functional connectivity (dFC) was estimated for each subject and trial using a sliding window. Phase Locking Value (PLV) was used to quantify the statistical coupling between ROIs. Finally, the temporal Independent Component Analysis (tICA) was applied on the dFC tensor to extract dynamic brain network states, including spatial network maps and temporal activity. A null distribution was generated to assess the temporal moments of significant modulation for each of the extracted states (as highlighted in red).

### 2.4 EEG Source Connectivity

#### i. Forward Model

Following the equivalent current dipole model, EEG signals measured from Q channels can be expressed as linear combinations of P time-varying current dipole sources as follows:

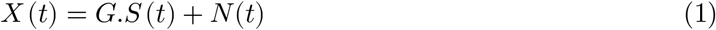

Where *G*(*Q*×*P*) represents the forward model, often called the lead field matrix, and *N* (*t*) denotes the additive noise. The lead field matrix is computed from a realistic head model along with the position of electrodes. Here, we used the Boundary Element Method (BEM) head model fitted to the ICBM MRI template (Kötter et al., 2001), downloaded from https://www.mcgill.ca/bic/software/tools-data-analysis/anatomical-mri/atlases/icbm152lin using the OpenMEEG toolbox (Gramfort et al., 2010), and used the Electrical Geodesic Inc (EGI) configuration for the EEG electrodes.

#### ii. Inverse Solution: wMNE

The EEG inverse problem consists in estimating the unknown parameters of dipolar source S(t) at the cortical level (position, orientation, and magnitude), from the measured EEG signals X(t) at the scalp level. Here, we used the Destrieux atlas parcellation (148 ROIs) (Destrieux et al., 2010) to locate cortical sources, and constrained their orientation normally to the cortical surface (Dale and Sereno, 1993). Therefore, the EEG inverse problem was reduced to the estimation of sources magnitude:

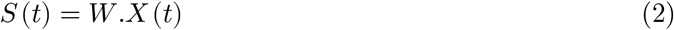

To compute the inverse matrix W, we used the weighted minimum norm estimate (wMNE) (Lin et al., 2006) method, that compensates the tendency of the classical minimum norm estimate (MNE) (Hämäläinen and Ilmoniemi, 1994) to favor weak and surface sources:

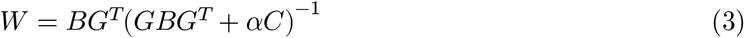

where *B* is the diagonal weighting matrix (inversely proportional to the norm of lead field vectors), is the lead field matrix, *α* is the regularization parameter (based on signal to noise ratio: *α* = 1/*SNR*), and *C* is the noise covariance matrix (calculated from our 700 ms pre-stimulus baseline). Here, we used the Matlab function implemented in the Brainstorm toolbox to compute wMNE. The SNR was set to 3, and the depth weighting value to 0.5 (default values).

#### iii. Dynamic Functional Connectivity: PLV - sliding window

Several approaches have been proposed to compute functional connectivity between reconstructed regional time series. In this study, we used the phase-locking value (PLV) method (Lachaux et al., 2000). Since we aimed to assess functional connectivity dynamics, we chose the sliding window approach defined by its length with an overlapping step. Hence, for each correct incongruent trial, PLV measures the phase synchronization between two signals *x*(*t*) and *y*(*t*) within each temporal window t:

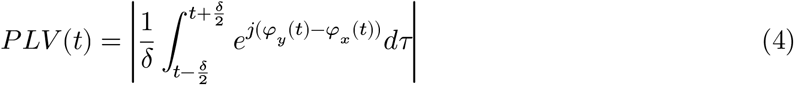

where *φ*_*y*_ (*t*) and *φ*_*x*_ (*t*) are the instantaneous phases of signals *y*(*t*) and *x*(*t*), respectively derived from the Hilbert transform.

The choice of temporal window length was based on (Lachaux et al., 2000), where it is recommended to consider at least 6 cycles at the frequency band of interest as a compromise between temporal and spatial accuracy. In this study, we conducted our analysis separately in both the beta [12-25Hz] (central frequency: *Cf*= 18.5 Hz) and gamma bands [30-45Hz] (central frequency: *Cf*= 37.5 Hz). Thus, we chose the smallest window length equals to 6/Cf, that is 320ms for the beta band and 160ms for the gamma band. We considered a 90% overlap between consecutive windows to track fast neural activity. Therefore, the total number of windows over the whole trial duration was *nWinds* = 49 windows for the beta band, and nWinds = 109 windows for the gamma band, and the output dimension of the dynamic functional connectivity (dFC) matrix was *nROIS* × *nROIS* × *nWinds* for each subject trial.

### 2.5 Dynamic Brain Network States (dBNS)

#### i. Temporal Independent Component Analysis (tICA): JADE

For each subject’s correct incongruent trial, the dFC tensor can be unfolded into a 2D matrix of dimension *nROIS*(*nROIS* − 1)/2 × *nWinds* due to symmetry. Then, for each group separately, the resultant dFC matrices of all trials and subjects were concatenated along the temporal dimension to generate a group-specific dFC matrix denoted *M*. *M*_*C*_ refers to the dFC relative to the HC group, while *M*_*P*_ refers to the dFC relative to the PD group. Since we aimed to summarize and extract the most relevant time-varying connectivity patterns in both MC and MP, this problem can be formulated as follows:

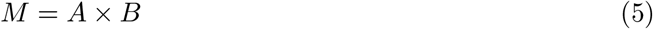

Where *A* is the mixing matrix that represents the spatial maps of dominant brain network states, and B describes their temporal evolution. Among the existing decomposition and clustering techniques, we derived the dynamic Brain Network States (dBNS) using temporal Independent Component Analysis (tICA) adopted by several previous studies (O’Neill et al., 2017; Yaesoubi et al., 2015). This technique assumes maximal independence between the time courses of the extracted dBNS. Here, tICA was performed using the JADE algorithm (Joint Approximate Diagonalization of Eigen-matrices) (Cardoso and Souloumiac, 1993; Rutledge and Jouan-Rimbaud Bouveresse, 2013). Briefly, JADE applies the Jacobi technique to optimize contrast functions based on high statistical order (Fourth Order: FO) cumulants of the data. One advantage of JADE compared to other ICA methods (FastICA, infomax, …), in our situation, is its high robustness for small sample sizes: Tabbal et al., 2021 showed that group-subject similarity was above 90% when there were only 5 subjects, and 100% with as few as 7 subjects. Here, the JADE function in Matlab was used (The Mathworks, USA, version 2019a).

#### ii. Number of states selection

Determining the optimal number of states to be extracted by tICA is a crucial issue for most decomposition and dimensionality reduction methods (Cong et al., 2013; Mørup and Hansen, 2009). Here, we used the DIFFIT (difference in data fitting) method based on the goodness-of-fit approach (Timmerman and Kiers, 2000; Wang et al., 2018), previously used by recent studies (Tabbal et al., 2021; Tewarie et al., 2019; Zhu et al., 2020). DIFFIT is calculated based on the following equations:

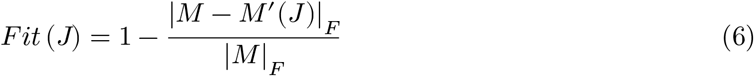

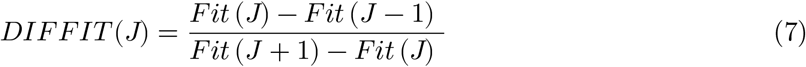

Where *M* is the original concatenated dFC matrix to be decomposed, and *M*′ is the reconstructed matrix after tICA decomposition at the number of states *J*, and || is the Frobenius norm. We varied *J* from 3 to 10 states, then chose the number of states *J* resulting in the largest DIFFIT value. For each group, we applied the DIFFIT method on the dFC matrix of each subject separately, then averaged the obtained DIFFIT values across subjects to obtain an average of 5 dynamic brain network states (dBNS) for both HC and PD groups (mean=5.3 for HC, mean=4.9 for PD).

#### iii. Significant task-modulated states

An additional step was followed to automatically select, from all extracted dBNS, those that were significantly modulated by the task for each group. First, we built an empirical null distribution through the generation of a surrogate time course based on a sign-flipping permutation following the procedure detailed in previous studies (Hunt et al., 2012; O’Neill et al., 2017; Tabbal et al., 2021; Winkler et al., 2014). A dBNS was considered as significant if its corresponding time course fell outside the distribution for a duration determined by at least 3 successive cycles (a cycle was determined based on the lower band of the studied frequency interval). 2-tailed distribution was allowed, followed by a Bonferroni correction accounting for multiple comparisons across the extracted dBNS (see (O’Neill et al., 2018, 2017) for details). Consequently, a set of significant states for the HC group, and significant states for the PD group were obtained.

### 2.6 Between-Group Statistical Differences

Finally, we intended to quantify the statistical differences between HC and PD groups regarding the dBNS defined by tICA. To achieve this, we used a back-fitting approach applied to EEG microstates, previously used at the sensor-level (Khanna et al., 2015; Michel and Koenig, 2018), as described below and illustrated in Figure 3.

**Figure 3.**
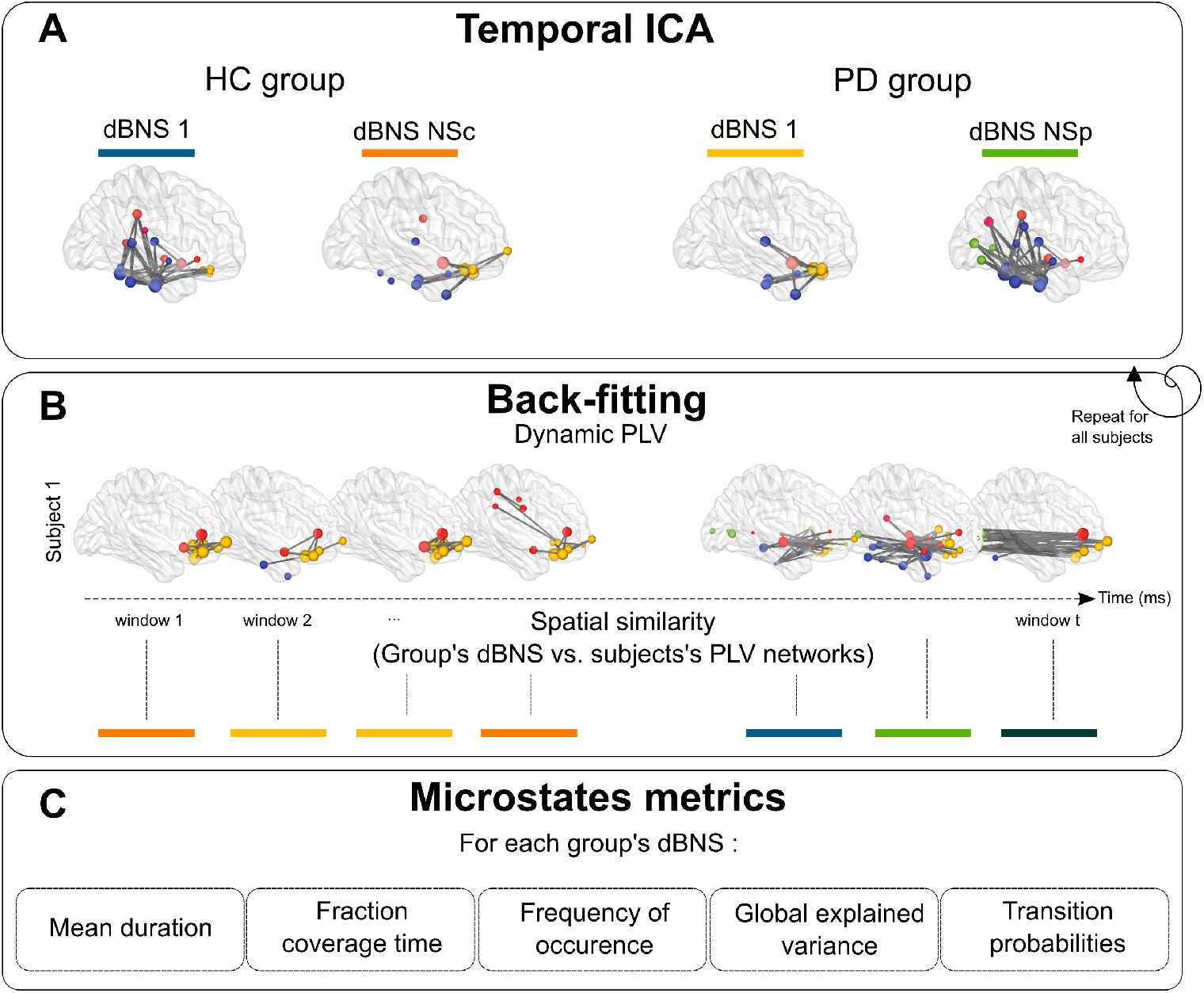
A. Temporal ICA applied to HC and PD groups. The resultant significant and ICA networks for HC and PD groups respectively are illustrated by specific colors. B. Back-Fitting approach assigns PLV network at each temporal window with one of all (PD and HC) the ICA networks (microstate) having the highest spatial similarity value. This was applied on all HC and PD subjects, followed by a calculation of the main network states metrics as shown in C.

#### i. Back-Fitting

First, all selected and states were assigned separately to all participants (HC and PD patients) and to every temporal window of all individual dFC, using the approach proposed by (Ville et al., 2010) in the context of classical scalp-level ‘microstates’. The spatial similarity (correlation) was calculated between the dFC map at every temporal window and each of the dBNS. Then, using a ‘winner-takes-all’ approach, each temporal window was labeled with the best-fitting dBNS (with the highest spatial similarity). Therefore, we obtained a temporal dBNS sequence for each individual dFC for both groups.

#### ii. Network States Metrics

Several features of the various dBNS, originally coming from the microstate literature, can be computed separately for each HC and PD subjects (Lehmann et al., 2005). These metrics will be referred to as network states metrics for the rest of the paper in order to avoid potential confusions in interpretation. Our features of interest were:

- Average lifespan (or mean duration; in seconds). The lifespan of a dBNS was calculated as the average time during which a given dBNS remains stable for successive segments (Lehmann et al., 1987).
- Fraction coverage time (in percentage between 0 and 1). The coverage is defined as the ratio of the time frames for which a given dBNS is dominant relative to the total recording duration (Lehmann et al., 1987).
- Frequency of occurrence (in Hz). The frequency of occurrence represents the number of unique appearances of the dBNS per second, independently of its duration (Lehmann et al., 1987).
- Global Explained Variance (GEV; in percentage between 0 and 1). The GEV of a dBNS is the percentage of the total variance explained by this dBNS (Brodbeck et al., 2012).
- Transition probabilities. The transition probabilities are defined as the sequence of transitions from one dBNS to another (Lehmann et al., 2005).

### 2.7 Statistical Analysis

All statistical analyses were performed using R version 4.0.2 (R Core Team, 2020) implemented with the lme4 package for mixed model analyses (Bates et al., 2015).

#### i. Behavior

RT and accuracy are the variables measured during the Simon task. Comparing congruent and incongruent RT of correct responses provides an estimate of the congruence effect that informs about the additional time needed to solve conflict. Incongruent trials are typically associated with increased RT and decreased accuracy. In addition to the analysis of the congruence effect, behavioral data were also analyzed in light of the activation-suppression model (Ridderinkhof, 2002), which informs about the temporal characteristics of cognitive action control. These are based on distributional analyses that measure impulsive action selection (incongruent accuracy for the fastest trials) and suppression (slope value of the congruence effect for the slowest trials). A complete description of the model and analysis steps can be found in van den Wildenberg et al. (2010) and in Duprez et al. (2017). Briefly, RT are increasingly ordered and divided into 7 bins. Accuracy and congruence effect (incongruent RT - congruent RT) are then calculated for those 7 bins resulting in two different representations : conditional accuracy functions and delta plots, respectively. The activation-suppression model postulates that incongruent accuracy of the first bin (fastest trials) informs about impulsive action selection, while the slope between the two last bins of the delta plots informs about selective suppression of such impulsive actions.

The effect of congruence and group on RT and accuracy were first broadly analysed by using linear and non-linear mixed models, respectively. In both cases, congruence and group were added as fixed factors, while a random slope and intercept was allowed for each subject. RT were log-transformed for increased compliance with the model’s assumptions in that case. The formulas used for the models were the following:

RT model :

Model = lmer(log(RT)∼condition × group + (condition | subject), data =data)

Accuracy model :

Model = glmer(accuracy∼condition × group + (condition | subject), family = binomial, glmerControl(optimizer = “bobyqa”), data =data)

Fixed effects significance were computed through the Anova function of the {car} package (Fox and Weisberg, 2019) that calculates type II Wald chi-square tests. Marginal (mR^2^) and conditional (cR^2^) R^2^ were calculated using the {MuMin} package (Barton, 2009).

Average first bin incongruent accuracy and last slope values of conditional accuracy and delta plots were directly extracted from subjects in both groups. Given the unequal sample sizes between the groups (10 HC vs 21 PD), comparisons were performed using Welch’s t test and are reported with effect size (Cohen’s d).

#### ii. Network states metrics

To quantify the differences between HC and PD groups in terms of network states parameters (mean duration, fraction coverage time, frequency of occurrence, GEV, and transition probabilities), statistical tests were performed at the level of each extracted group ICA component. Welch’s t-test was used here also to assess the statistical difference between the two groups. Note that, given that each ICA component is independent, no corrections were applied for multiple testing over the different dBNS.

### 2.8. Code availability

All the Matlab and R codes used for source reconstruction, dFC, tICA, backfitting and microstate metrics, as well as all subsequent statistical analyses are publicly available at https://github.com/judytabbal/dynCogPD.

## 3. Results

This section first presents the behavioral results. Then, the EEG network results are divided in group-level results, presenting the significant dBNS derived from tICA, and subject-level results, relative to each subjects’ network states metrics of significant dBNS.

**Table 1:**
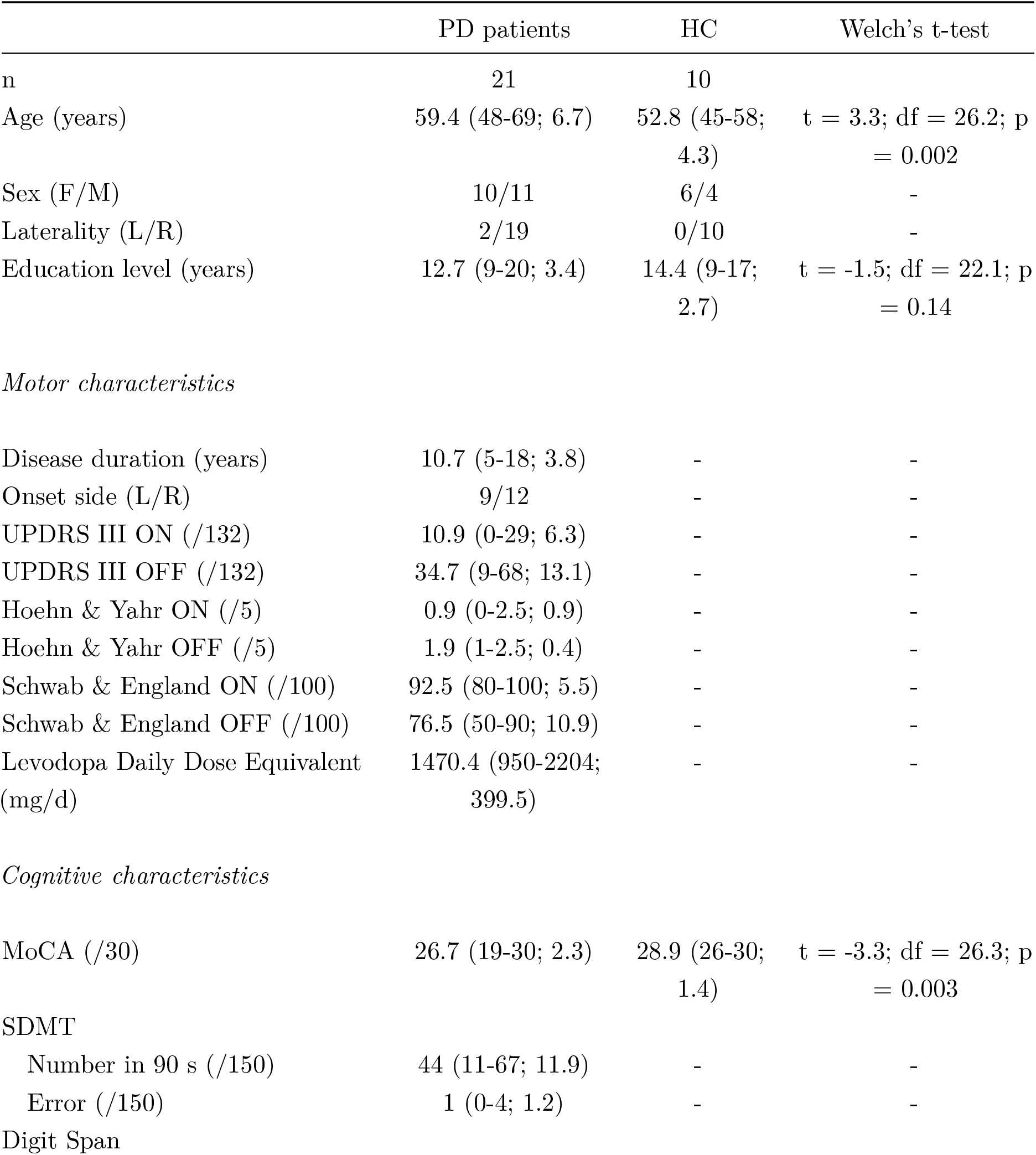
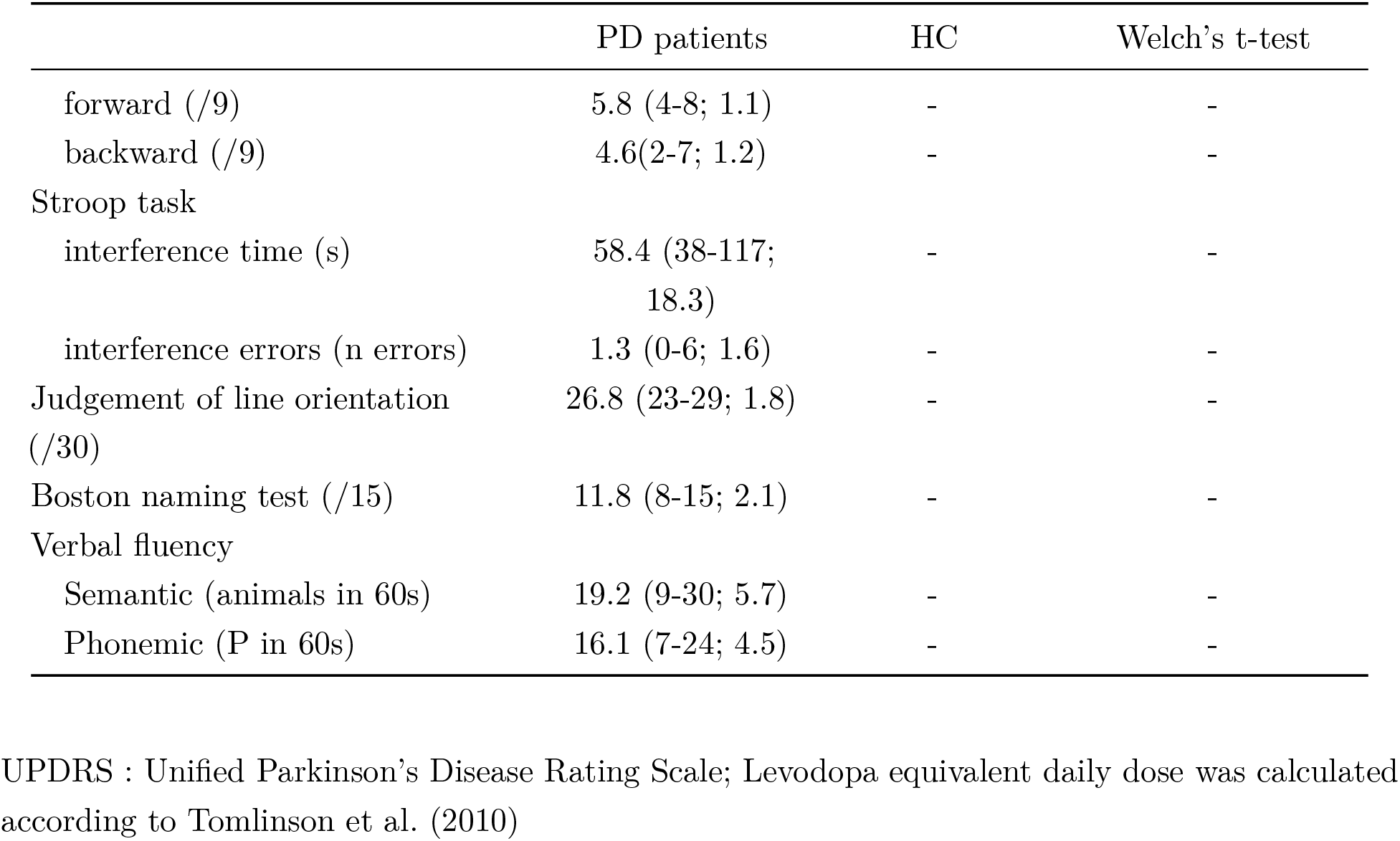
demographic and clinical data. Data are presented as mean (range; sd)

|  | PD patients | HC | Welch's t-test |
| --- | --- | --- | --- |
| n | 21 | 10 |  |
| Age (years) | 59.4 (48-69; 6.7) | 52.8 (45-58; 4.3) | t = 3.3; df = 26.2; p = 0.002 |
| Sex (F/M) | 10/11 | 6/4 | - |
| Laterality (L/R) | 2/19 | 0/10 | - |
| Education level (years) | 12.7 (9-20; 3.4) | 14.4 (9-17; 2.7) | t = -1.5; df = 22.1; p = 0.14 |
| *Motor characteristics* |  |  |  |
| Disease duration (years) | 10.7 (5-18; 3.8) | - | - |
| Onset side (L/R) | 9/12 | - | - |
| UPDRS III ON (/132) | 10.9 (0-29; 6.3) | - | - |
| UPDRS III OFF (/132) | 34.7 (9-68; 13.1) | - | - |
| Hoehn & Yahr ON (/5) | 0.9 (0-2.5; 0.9) | - | - |
| Hoehn & Yahr OFF (/5) | 1.9 (1-2.5; 0.4) | - | - |
| Schwab & England ON (/100) | 92.5 (80-100; 5.5) | - | - |
| Schwab & England OFF (/100) | 76.5 (50-90; 10.9) | - | - |
| Levodopa Daily Dose Equivalent (mg/d) | 1470.4 (950-2204; 399.5) | - | - |
| *Cognitive characteristics* |  |  |  |
| MoCA (/30) | 26.7 (19-30; 2.3) | 28.9 (26-30; 1.4) | t = -3.3; df = 26.3; p = 0.003 |
| SDMT |  |  |  |
| Number in 90 s (/150) | 44 (11-67; 11.9) | - | - |
| Error (/150) | 1 (0-4; 1.2) | - | - |
| Digit Span |  |  |  |
| forward (/9) | 5.8 (4-8; 1.1) | - | - |
| backward (/9) | 4.6(2-7; 1.2) | - | - |
| Stroop task |  |  |  |
| interference time (s) | 58.4 (38-117;<br>18.3) | - | - |
| interference errors (n errors) | 1.3 (0-6; 1.6) | - | - |
| Judgement of line orientation<br>(/30) | 26.8 (23-29; 1.8) | - | - |
| Boston naming test (/15) | 11.8 (8-15; 2.1) | - | - |
| Verbal fluency |  |  |  |
| Semantic (animals in 60s) | 19.2 (9-30; 5.7) | - | - |
| Phonemic (P in 60s) | 16.1 (7-24; 4.5) | - | - |

UPDRS : Unified Parkinson's Disease Rating Scale; Levodopa equivalent daily dose was calculated according to Tomlinson et al. (2010)

### 3.1. Behavioral results

The typical congruence effect was found, showing increased RT (*χ*^2^ = 196, p<0.0001; mR^2^ = 0.11; cR^2^ = 0.50) and decreased accuracy (*χ*^2^ = 76.2, p<0.0001; mR^2^ = 0.51; cR^2^ = 0.28) in the incongruent condition (Figure 4A and B). PD patients were slower than HC regardless of congruence (*χ*^2^ = 5.6, p = 0.018), but were overall similarly accurate (*χ*^2^ = 0.74, p = 0.39). The congruence effect was not significantly different between groups, both for RT (*χ*^2^ = 0.06, p = 0.80) and accuracy (*χ*^2^ = 0.25, p = 0.62). Conditional accuracy functions revealed the classic pattern of increased accuracy with RT (see supplementary figure SF1A-B). Accuracy of the first bin in the incongruent condition reflects impulsive action selection (Figure 4C), and did not significantly differ between PD patients and HC (t = 0.21, p = 0.84). Inspection of delta plots (see supplementary Figure SF1C) also exhibited the typical decreasing pattern of the congruence effect associated with the Simon task. The last slope of delta plots informs on the strength of selective inhibition of inappropriate responses (Figure 4D). In line with our hypothesis, PD patients had a significantly flatter slope, indicative of reduced proficiency in inhibiting automatic responses (t = -2.64, p = 0.02; Cohens’ d = 1.29).

**Figure 4.**
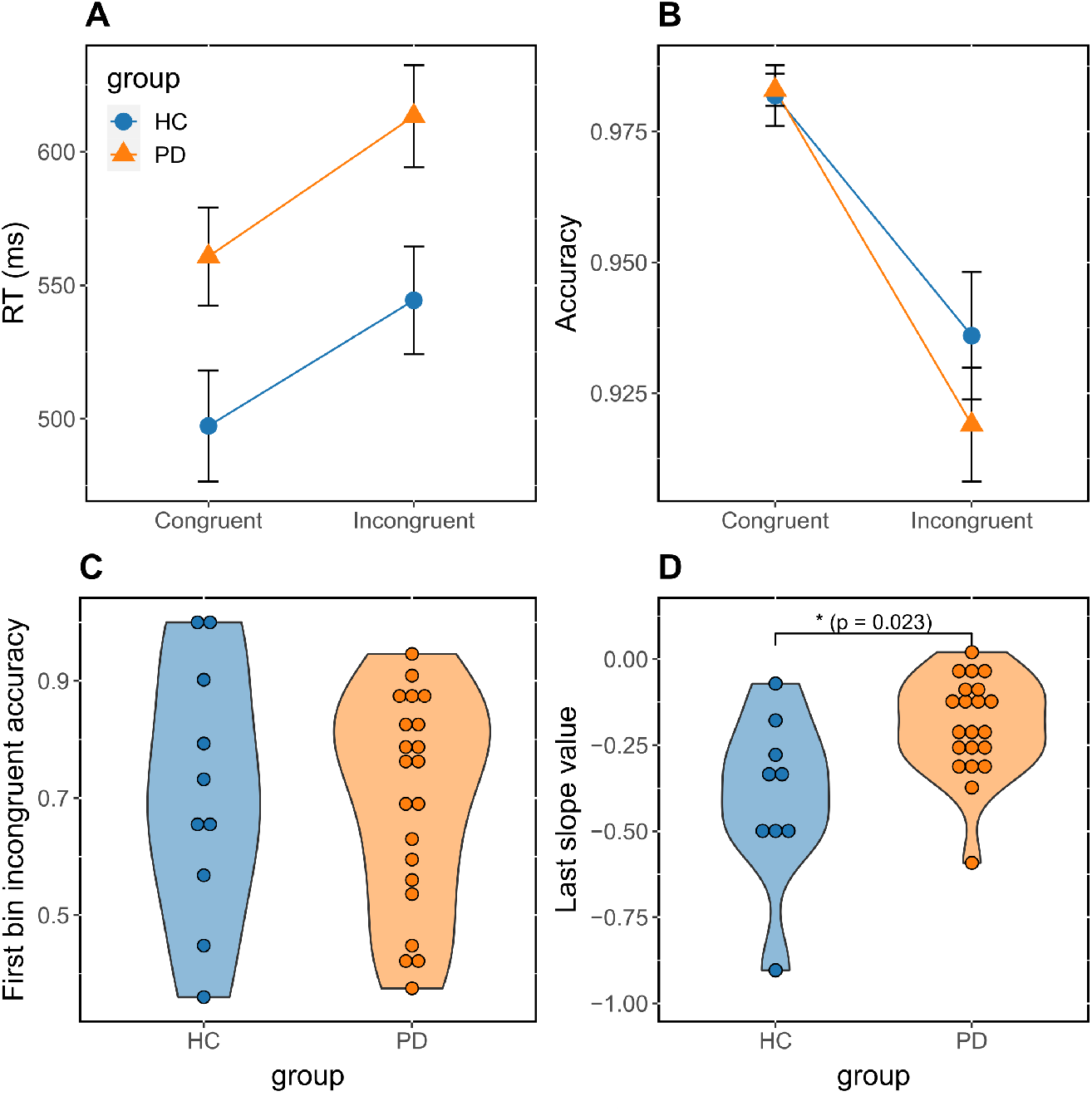
Average RT (A) and accuracy (B) as a function of congruence in both groups. Error bars represent standard error of the mean. Impulsive action selection is denoted by group-specific violin plots showing accuracy of the first incongruent bin (C) of conditional accuracy function. The strength of selective inhibition is represented by group-specific violin plots showing the last slope value (D) of delta plots.

### 3.2. Group-level results – Significant task-modulated dBNS

In this section, we focus on the dynamics of the brain networks derived at the group-level from tICA. In the following, C_i_ denotes the dBNS_i_ extracted from the HC group, with i= [1; NSc], and P_j_ the dBNS_j_ from the PD group, with j= [1; NS_p_]. Only the dBNS that were significant following permutation testing and during at least a minimum of 3 cycles are presented here. However, all tICA-derived dBNS (significant or not) and their dynamic modulation (along with the null distribution) can be found in Supplementary Figures SF2-3. Our results show that, in the beta band, one significant dBNS was found in the HC group (NS_c_ = 1), and two in the PD group (NS_p_ = 2). In the gamma band, two significant dBNS were derived from the HC group (NS_c_ = 2), and three from the PD group (NS_p_ = 3). The spatiotemporal dynamics of beta and gamma dBNS are illustrated in Figure 5 and Figure 6, respectively. Since we aimed at tracking the evolution of the task-related components, we have plotted on the same time axis each group-specific dBNS and marked the corresponding duration for which they had a significant modulation. For the sake of visualization, we plotted the top 0.5% edges relative to the total number of unique possible connections, i.e., the top 55 edges per network. Using this threshold, in order to globally describe the integrated brain regions and characterize functional networks, we calculated the percentage of nodes with respect to the total number of ROIs, for each of the five macroscopic regions (F: Frontal, T: Temporal, P: Parietal, O: Occipital, and C: Cingulate/insula). In the description of the networks, only some of the nodes involved in the networks are discussed for clarity and are summarized based on larger regions they belong to. However, the reader can refer to Supplementary Table ST1 for a complete description of the Destrieux atlas regions involved in the networks, following the thresholding we previously described.

**Figure 5.**
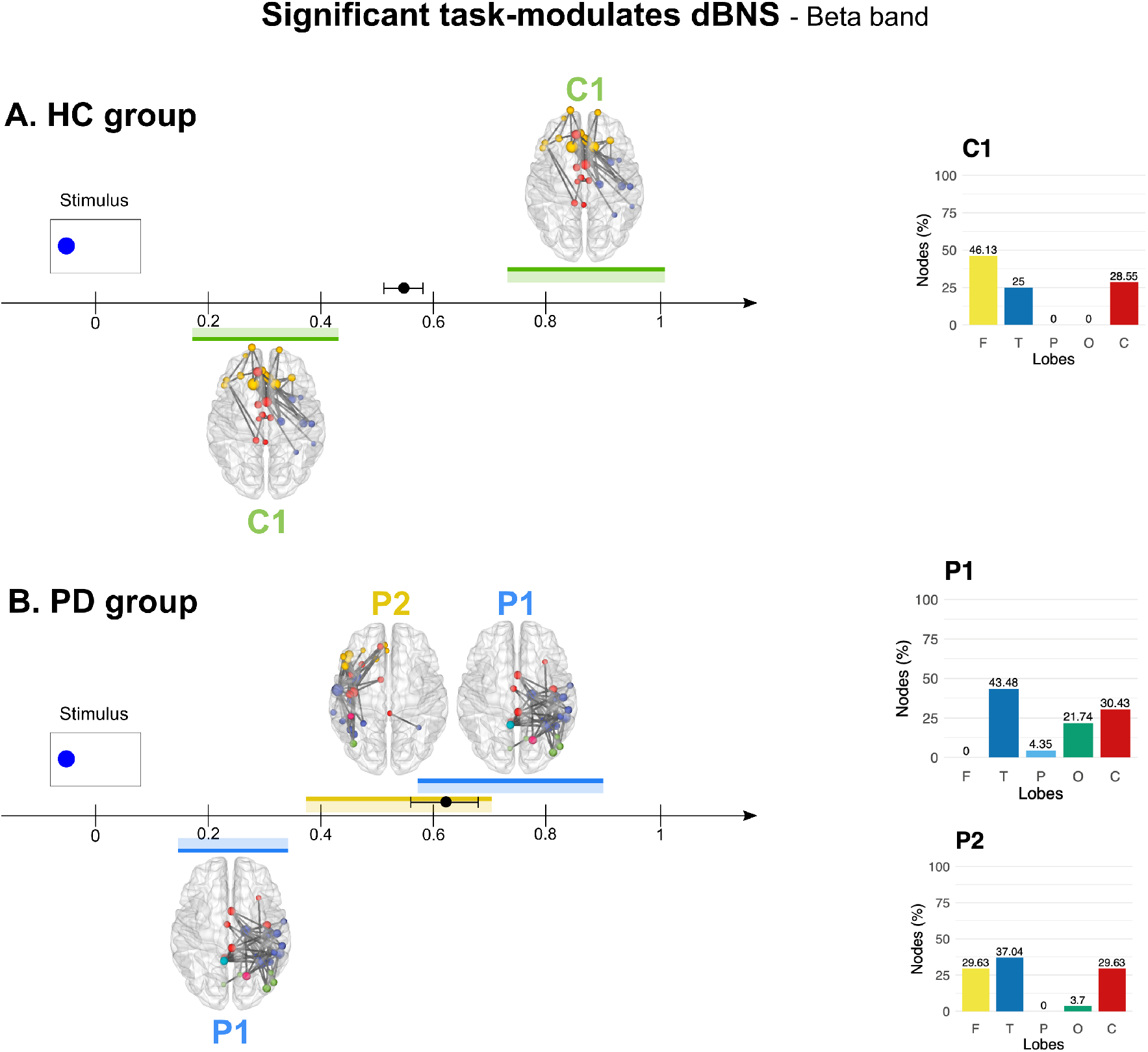
Spatiotemporal dynamics of the significant task-modulated dBNS for both HC (A) and PD (B) groups in the beta band. Time 0 s corresponds to stimulus onset. Each significant dBNS is illustrated as a brain network with a specific color. All brain networks were thresholded (only the top 0.5% edges are shown) for visualization. Node size is proportional to the degree (number of edges incident to a node). A color code is attributed for all nodes belonging to the same brain lobe (yellow for frontal, blue for temporal, light blue for parietal, green for occipital, and red for cingulate and insula). For each dBNS, we indicated the temporal duration during which it is significantly modulated by the task (positively modulated are plotted above the time axis, negatively modulated are plotted below the time axis). Overlaid data point in black corresponds to the average correct incongruent RT ± sd (of all trials). On the right side, the percentages of nodes relative to each brain lobe are illustrated on the colored bars for each state. The reader can refer to Supplementary Figure SF2 for a detailed representation of the network (top, left, and right views) with the corresponding averaged-trial temporal signals plotted over the whole temporal duration along with the null distribution to reveal the temporal significance. In supplementary table ST1, the labels of the activated Destrieux ROIs are highlighted.

**Figure 6.**
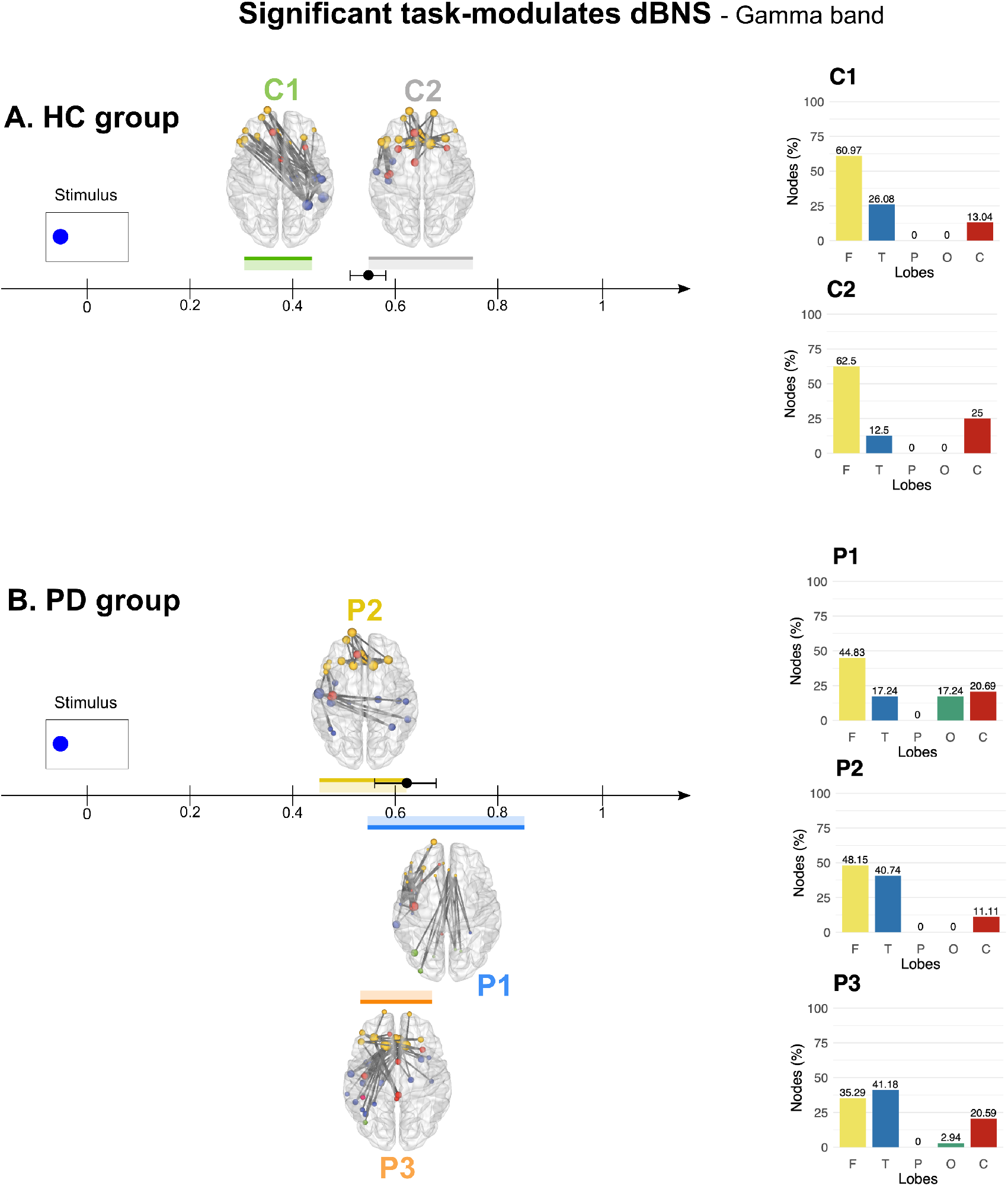
Spatiotemporal dynamics of the significant task-modulated dBNS for both HC (A.) and PD (B.) groups in the gamma band. The time 0 s corresponds to the stimulus offset. Each significant dBNS is illustrated as a brain network with a specific color. All brain networks were thresholded for visualization. Spheres of different sizes proportional to their strength represent the activated brain nodes. A color code is attributed for all nodes belonging to the same brain lobe (yellow for frontal, blue for temporal, light blue for parietal, green for occipital, and red for cingulate and insula). For each state, we indicated the temporal duration on which it is significantly modulated by the task (positively modulated are plotted above the time axis, negatively modulated are plotted below the time axis). Overlaid data point in black corresponds to the average correct incongruent RT ± sd (of all trials). On the right side, the percentages of nodes relative to each brain lobe are illustrated on the colored bars for each state. The reader can refer to Supplementary Figure SF3 for a detailed view representation of the network (top, left, and right) with the corresponding averaged-trial temporal signals plotted over the whole temporal duration along with the null distribution to reveal the temporal significance. IIn supplementary table ST1, the labels of the activated Destrieux ROIs are highlighted.

#### 3.2.1. Beta band

In the beta band, C1 (Figure 5A) was significantly modulated by the task in the HC group during two different periods, before and after the group-averaged RT (544 ms, sd = 64), ranging from 130 to 460 ms, with a drop in connectivity; then from 700 to 1000 ms showing increased connectivity. This network mainly involves connections in the (i) frontal lobe (46%) bilaterally, including frontopolar, orbital and inferior frontal regions, (ii) the right temporal lobe (25%), including the temporal pole and parahippocampal and fusiform regions, and (iii) the cingulate lobe (29%) bilaterally, with anterior/posterior cingulate and pericallosal regions.

Regarding the PD group, a dBNS P1 was found to be significantly modulated by the task with a temporal variation similar to C1 (desynchronization between 130 and 360 ms, followed by synchronization between 580 and 870 ms, overlapping with averaged RT : 613 ms, sd = 87). However, the two networks differed spatially, notably in the absence of frontal nodes in P1. In P1, functional connectivity was mostly present in the right hemisphere and dominated in the temporal lobe (43%), including superior and inferior temporal as well as fusiform regions. P1 is also characterized by deeper connections (30%), with posterior cingulate, sub/pericallosal and insular regions, and also by occipital connections (22%), including inferior occipital and occipito-temporal regions. Finally, P1 also has connections in the parietal lobe, but to a lesser extent (5%), with only subparietal regions.

Another dBNS denoted P2 was significantly modulated by the task in the PD group with increased connectivity from 0.39 to 0.71 s, overlapping with averaged RT. In this network, connections were mostly distributed over the left hemisphere. Temporal nodes dominated (38%) and included the temporal pole, fusiform area, as well as superior, middle and inferior temporal regions. Frontal connections were also significant (29%) and were represented by inferior frontal, orbital and rectus regions. Deeper connections were equally present (29%), with anterior and posterior cingulate cortices, as well as insular regions. Occipital lobe connections were also present but to a lesser extent (4%), and involved occipito-temporal (lingual gyrus) regions.

As a whole, we found that PD patients and HC displayed spatially different dBNS after presentation of an incongruent stimulus with a beta dBNS involving frontal, cingular, and temporal connections in HC that was completely absent in PD. Instead, dBNS with lateralized temporo-insular-occipital and temporo-frontal connections were found in PD that were absent in HC.

#### 3.2.2. Gamma band

In the gamma band, two dBNS were identified in the HC group (Figure 6). First, C1 was significantly modulated by the task with increased connectivity before the averaged RT from 0.32 to 0.42 s. C1 consisted mostly in frontal nodes (61%), predominantly in the left hemisphere, including inferior frontal, frontopolar, frontomarginal and orbital regions. Frontal nodes were highly connected with right temporal areas (26%) represented by superior and inferior temporal areas, as well as fusiform and parahippocampal regions. C1 also included deeper nodes (13%) including the left anterior cingulate and right anterior insula.

A second dBNS, C2, was significantly modulated by the task and had increased connectivity beginning around averaged RT, from 0.56 to 0.72 s. As in C1, there was a high contribution of frontal nodes (62%) but bilaterally; while C1 showed mostly left frontal nodes. These included inferior frontal, frontopolar, frontomarginal, rectus and orbital regions. Contrary to C1, C2 did not include a left frontal-right temporal connectivity pattern. Temporal nodes were less prominent (13%), and were restricted to the left hemisphere, including superior temporal and anterior collateral sulcus regions. Deep nodes were more present (25%), and were found mostly in the left hemisphere, with anterior cingulate, subcallosal, and anterior, central and inferior insular regions.

Regarding the PD group, three dBNS (P1, P2, and P3) were significantly modulated by the task. These 3 dBNS followed each other with overlapping periods in that order: P2 showing increased connectivity from 0.44 to 0.61 s; then P3, with decreased connectivity from 0.53 to 0.68 s; and finally P1, exhibiting decreased connectivity from 0.58 to 0.84 s. Significant modulation of P2 terminated around the averaged RT, while P1 started around RT, and the significant modulation period of P3 overlapped entirely with averaged RT.

P1 consisted in left fronto-temporal and fronto-insular connections, as well as long-range bilateral fronto-occipital connections. Frontal nodes dominated (45%), especially in the left hemisphere, including inferior frontal, frontomarginal, frontopolar and orbital regions. Temporal regions (17%) were represented by superior and middle temporal areas, while the occipital lobe, which was equally represented (17%), included calcarine, anterior occipital and occipito-temporal areas. Finally, deep structures (21%), only represented in the left hemisphere, involved the anterior and posterior cingulate, as well as anterior, central and inferior insular regions.

P2 was characterized mostly by left-right temporo-temporal and fronto-frontal connections. Frontal regions (48%) involved orbital and rectus regions, as well as frontopolar, frontomarginal and inferior frontal areas. Temporal nodes (41%) included fusiform and parahippocampal structures, as well as superior and inferior temporal areas. P2 did not include any occipital or parietal regions. However, deep structures were found (11%), only in the left hemisphere, and were represented by anterior cingulate, central and inferior insular regions.

P3 was characterized mostly by left fronto-temporal connections, as well as fronto-insular/cingulate and bilateral fronto-frontal connections. Frontal nodes (35%) involved mostly orbital, frontopolar and rectus regions. The temporal lobe (41%) was mostly represented by left hemisphere structures including inferior, middle and superior temporal areas, as well as the temporal pole and parahippocampal regions. Occipital nodes (3%) included only left occipito-temporal structures. Finally, deep structures (21%) involved subcallosal, anterior and posterior cingulate regions, and also anterior and inferior insular areas.

Overall, these results show again that incongruent trials in HC and PD patients modulate spatially distinct networks with various dynamics. While HC showed a sequential activity of two distinct dBNS involving fronto-temporal and fronto-cingular connectivity, PD patients had 3 overlapping dBNS with frontal, cross-hemispheric temporal as well as fronto-occipital connectivity.

### 3.3. Subject-level results - network states metrics analyses

As described in the Materials and Methods section, several network states metrics were computed for all subjects in both the beta and gamma bands, before performing the statistical analysis to quantify potential significant differences between groups. These are based on correlation between the dFC matrices with the dBNS, to determine, at any given time, which dBNS corresponds the most to the current network. Correlation between all dBNS (estimated separately in HC and PD) were performed with the dFC matrices of all subjects (e.g. correlations between an HC-defined dBNS was calculated with each HC subjects’ dFC matrices, and also with each PD subjects’ dFC matrices). The metrics computed thereafter included (i) average lifespan, (ii) frequency of occurrence, (iii) fraction coverage time, (iv) global explained variance, and (v) transition probabilities. These results are presented in Figure 7 for the beta band, and in Figure 8 for the gamma band.

**Figure 7.**
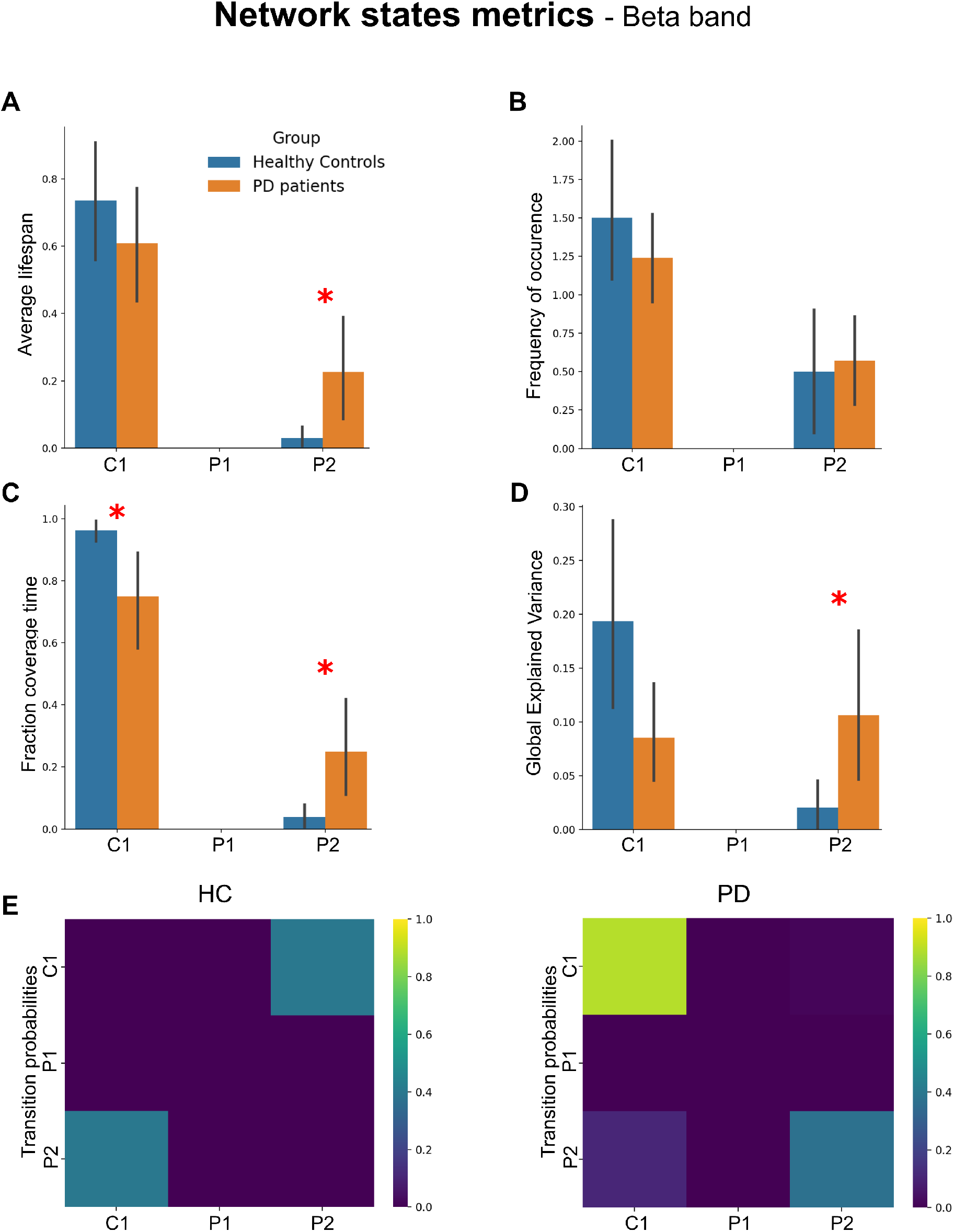
Network states parameters and statistical analysis between HC and PD groups in the beta band. The average lifespan, frequency of occurrence, fraction coverage time, and global explained variance results are represented by colored bars (mean value across subjects) (blue for HC group and orange for PD group) in A, B, C, and D. The standard deviations of the corresponding metrics are displayed as error bars. A red asterisk with the corresponding p-value illustrates the presence of statistical differences (Welsch’s Test; p-value<0.05) between the two groups in terms of microstate parameters. The transition probabilities between all dBNS are also shown in E.

**Figure 8.**
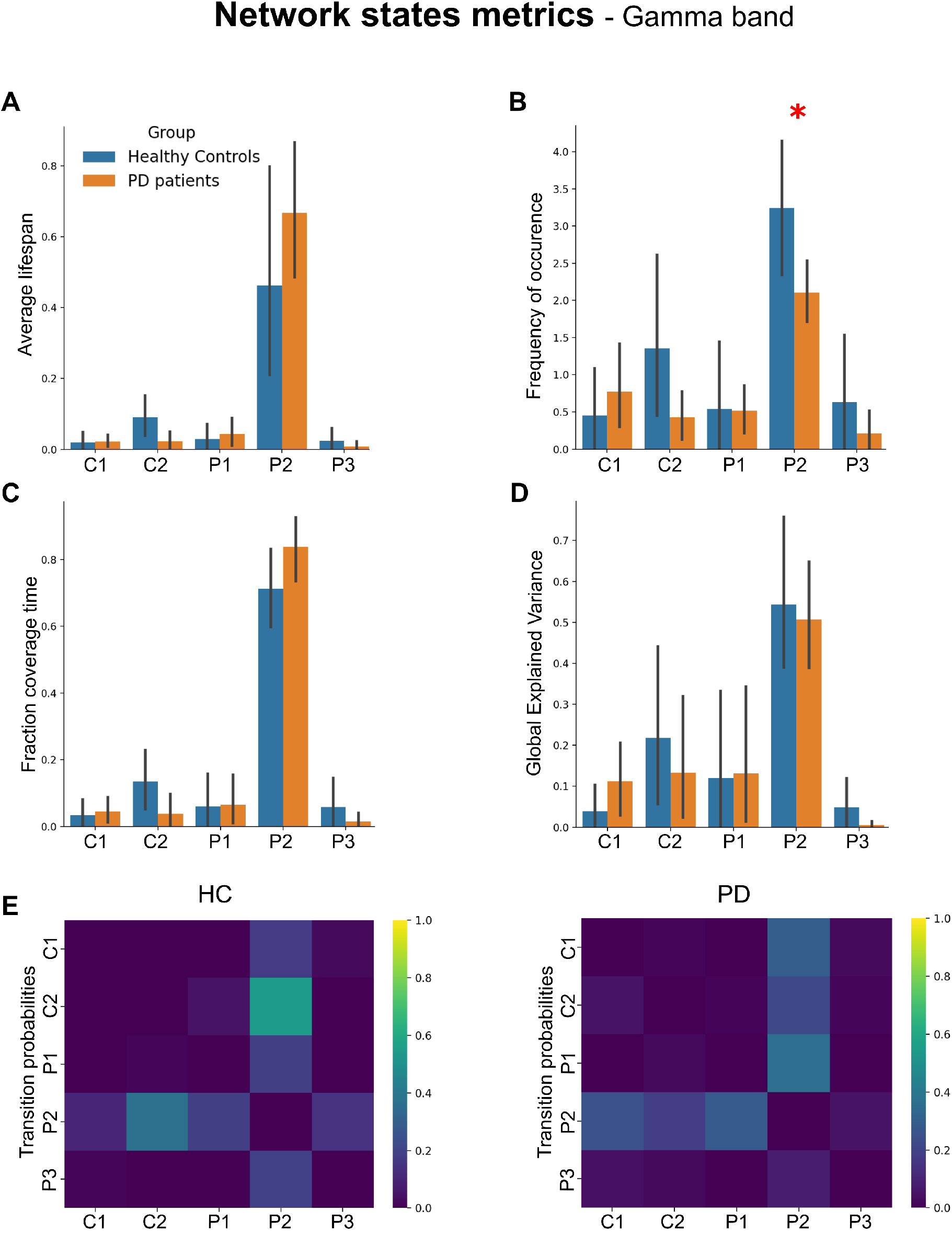
Network states parameters and statistical analysis between HC and PD groups in the gamma band. The average lifespan, frequency of occurrence, fraction coverage time, and global explained variance results are represented by colored bars (mean value across subjects) (blue for HC group and orange for PD group) in A, B, C, and D. The standard deviations of the corresponding metrics are displayed as error bars. A red asterisk with the corresponding p-value illustrates the presence of statistical differences (Welsch’s Test; p-value<0.05) between the two groups in terms of microstate parameters. The transition probabilities between all dBNS are also shown in E.

First, in the beta band, it can be noticed that all network states metrics relative to P1 were null (Figure 7). This is explained by the fact that, although P1 was significantly present at the group level (see previous section), this dBNS was never the dominating dBNS at the subject level (the dBNS that correlated the most with subjects’ dFC matrices), even in PD patients. Overall, all C1-related metrics seemed higher in HC than in PD. However, this was only supported by statistical analyses in the case of fraction coverage time (t = -2.52; p = 0.02; Cohen’s d = 0.78; all other comparisons : p>0.05). This result suggests that C1 was more present in HC than in PD, which is consistent with the fact that this dBNS was estimated based on HC data only. However, it is important to note that, although C1 was estimated on HC data, and although it seems more present in HC (based on a difference on one metric), it was the dominating dBNS in PD patients as well. Regarding P2, greater average lifespan (t = -2.45; p = 0.023; Cohen’s d = -0.76), fraction coverage time (t = -2.52; p = 0.02; Cohen’s d = -0.78), and global explained variance (t = -2.2; p = 0.036; Cohen’s d = -0.74) was found in PD patients as compared to HC. Frequency of occurrence did not differ between the two groups (t = -0.27, p = 0.79). These results indicate that, although present in HC, P2 was more present in PD patients. Finally, the transition probabilities results describe the sequential activity of the obtained dBNS (Figure 6E). Although dBNS of the HC group appeared to switch more often between C1 and P2 relative to the PD, this effect was not significant (t = 0.097, p = 0.92).

Regarding the gamma band, results of the comparison of network states metrics are displayed in Figure 8. Although all presented dBNS were significantly present at the group level (see section 3.2), the P2 dBNS largely dominated at the subject level in both groups. Importantly, although P2 was identified based on PD data, its frequency of occurrence was significantly higher in HC (t = 2.2, p = 0.045; Cohen’s d = 0.91). None of the other metrics regarding P2 had any significant differences between groups (all p > 0.05). All the other dBNS had similar metrics values between groups (all p > 0.05). From the transition probabilities results, Figure 8 suggests that HC had an increased transition probability between P2 and C2 as compared to PD patients. However, this difference was not significant (t = 1.8; p = 0.086). All other transition comparisons also revealed a similar transition probability between groups (all p > 0.05).

### 3.4. Correlation with behavioral/clinical data

We finally conducted correlations between behavioral, clinical data and EEG results. First, given that PD patients were older (t = 3.3; p = 0.002; Cohen’s d = 1.1) and had a lower MoCA score (t = -3.3, p = 0.003; Cohen’s d = -1.08) than HC, we calculated the correlation between age, MoCA and behavioral and EEG data. Age significantly correlated with congruent and incongruent RT (respectively : *ρ* = 0.55, p = 0.0015; *ρ* = 0.56, p = 0.0011; scatter plots can be found in supplementary figure SF4A-B), but with no other behavioral, clinical, or EEG variables (all p>0.05).

Given that behavioral group differences were restricted to RT and the last slope of the delta plots, we restricted our correlation analyses with EEG data to these measures and microstate metrics values of the significant dBNS that were present at the subject level. Consequently, dBNS microstate metrics that had 0 values, as that was the case for beta P1, could not be correlated with behavior and were thus ignored.

First regarding slope value, some dBNS, such as gamma C1 and gamma C2, were poorly represented at the subject level, with most subject values at 0 on network states metrics. Thus performing correlations with these networks’ metrics would lead to spurious findings.

The situation was different for the correlation between incongruent (and congruent) RT and the frequency of occurrence of beta C1, which was the most represented dBNS at the subject level in both HC and PD patients. Classic Spearman correlation could not be used, since frequency of occurrence took only 4 different discrete values in that case (0, 1, 2, or 3; see supplementary figure SF4C-D). Given that there were only 2 samples with a frequency of 0, and 2 with 3, we decided to ignore these values and computed a Welch’s t-test comparing congruent and incongruent RT between the frequency of occurrence values 1 and 2. We found that congruent and incongruent RT were higher when frequency of occurrence was lower (congruent RT: t = -2.95, p = 0.01, Cohen’s d = -1.19 ; incongruent RT: t = -2.7, p = 0.017, Cohen’s d = -1.08). In other words, slower RT were associated with a less frequent occurrence of beta C1.

## 4. Discussion

In this study, we aimed to explore i) how cognitive action control is associated with dynamically reconfiguring functional connectivity networks, and ii) how these dynamic structures are linked to PD-related alterations in CAC, as compared to healthy subjects. We used scalp HD-EEG recorded from 10 HC and 21 PD patients during a Simon task, and estimated source-reconstructed cortical functional networks within two frequency bands (beta and gamma). We applied a combination of wMNE with PLV (estimated through a sliding window approach) to track the dynamics of FC networks. To summarize dFC into a set of relevant connectivity patterns and to characterize their dynamics, a variant of temporal independent component analysis (tICA) was applied, providing a set of group-specific dynamic brain network (dBNS) states. The time-varying alterations in these states at the subject level were investigated using network states metrics. In both frequency bands, group-specific dBNS were found with specific spatial and temporal characteristics. Furthermore, microstate metrics analysis revealed group differences in the presence of some dBNS, especially in the beta band which were associated with reaction time at the task.

### 4.1. PD is associated with changes in cognitive action control

Among the various cognitive changes that have been associated with PD, cognitive action control (CAC) alterations are robustly reported. Some studies reported an increased global congruence effect (Praamstra et al., 1999, 1998; Schmiedt-Fehr et al., 2007; Wylie et al., 2005), others, which investigated the temporality of CAC, found changes in the dynamic expression of the process regarding impulsive action selection (Duprez et al., 2017; Wylie et al., 2009a) or suppression (van Wouwe et al., 2014; Wylie et al., 2010, 2009a, 2009b). In the present study, we found an overall slowing of PD patients as compared to HC, as well as a diminished strength of impulsive action suppression. Thus, our results are in line with the established description of CAC alterations in PD. It is important to note that we interpret the results according to the activation-suppression model (Ridderinkhof and others, 2002), and that other accounts of dynamic changes in accuracy and congruence effect have been proposed (for review, see Cespón et al., 2020).

### 4.2. Prefrontal beta is absent from PD dynamic brain networks during CAC

There has been a growing interest in the concept of functional brain states, which are characterized by a limited number of functional patterns with a temporarily stable activity followed by a fast transition to another state (Baker et al., 2014; Khanna et al., 2015; Michel and Koenig, 2018; O’Neill et al., 2018). Investigating functional brain states and their dynamic rearrangement implies the use of dimensionality reduction methods. Although K-means clustering has been commonly used in most aforementioned studies, other clustering/decomposition algorithms can also be applied to estimate dynamic brain states. Among these methods, we focused on temporal Independent Component Analysis (tICA), since it has proven its ability to track fast temporal variations in brain connectivity (O’Neill et al., 2017; Tabbal et al., 2021; Yaesoubi et al., 2015). tICA has also been applied for clinical purposes to characterize the spatiotemporal alterations induced in several brain disorders, as in Alzheimer’s Disease (Koelewijn et al., 2017), epilepsy (Koelewijn et al., 2015), and depression (Knyazev et al., 2016; Nugent et al., 2015) using EEG/MEG data during rest and task. Since CAC is a dynamic process, this approach could contribute to characterizing PD-related alterations of CAC in terms of involved brain networks.

Our results show that CAC in HC can be globally described by one dominant beta network comprising fronto-cingulo-temporal functional connections that decreased just before the correct response, and increased immediately after. Since we used a network-based approach, and because of the relatively high number of areas (nodes) involved in the network, it is challenging to envision a localizationist interpretation of those results. Nevertheless, it is worth noting that this network involves areas commonly associated with conflict resolution and inhibition found using fMRI, such as the right inferior frontal gyrus, the anterior cingulate cortex, as well as orbitofrontal cortex (Aron et al., 2004; Eagle et al., 2008; B. Forstmann et al., 2008; B. U. Forstmann et al., 2008; Widge et al., 2019; Yeung et al., 2007). Furthermore, prefrontal beta activity has been associated with cognitive control and attention (Friedman and Robbins, 2021; Schmidt et al., 2019; Swann et al., 2009 ; for review, see Engel and Fries, 2010; Wang, 2010). In addition, the dominant beta network in HC also involved temporal regions, including the fusiform area, which has been recently shown to be modulated by attentional demand in both humans and macaques (Kim et al., 2012; Sani et al., 2021; Wittfoth et al., 2006).

In addition to the nodes involved in this network, one could wonder about the general role of beta oscillations in such a network. In that respect, our results could be explained by a stimulus-induced desynchronization effect, followed by a beta rebound during the decision-making (as considered in event-related (de)synchronization studies) or as a spontaneous effect of the inhibition/activation process related to the conflict task (Panagiotaropoulos et al., 2013; Wu et al., 2019). However, our data showed that increased synchronization occured after the correct response, which excludes a decision-making interpretation. Another interesting account was proposed by Engel and Fries with the “Status Quo” hypothesis (Engel and Fries, 2010), which proposed that beta activity would be involved in sustaining a cognitive set when dealing with a task requiring a strong top-down component. This hypothesis is also hard to reconcile with our results, given that beta C1 showed decreased connectivity before the correct response, and increased connectivity after. Indeed, the period between incongruent stimulus onset and the correct response requires a strong top-down component. In light of Engel and Fries’ hypothesis, one would expect beta synchronization to be stronger in this period, and not weaker as we observed. That being said, cortical beta oscillations unlikely have a monolithic role, let alone in a task such as the Simon task. Indeed, the Simon task measures CAC, which in itself involves several other overlapping cognitive processes such as attention and working memory, or inhibition, and which involve strong response monitoring mechanisms. In that respect, beta oscillations were also associated with several other roles, such as activation of a cognitive set (Spitzer and Haegens, 2017), time estimation (Kulashekhar et al., 2016), default state of working memory (Lundqvist et al., 2016), or a clear-out of working memory after completion of a trial (Schmidt et al., 2019).

Interestingly, the strong bilateral frontal component that we observed in beta C1 was completely absent in one dBNS of PD patients. We found two independent dBNS in PD patients that were strongly lateralized. The first one (beta P1) was right-sided and mostly involved superior and inferior temporal regions, as well as cingulate and insular cortex, and inferior occipital areas. In beta P1, all the frontal regions usually associated with CAC were absent. The time-variation of this network is noteworthy given that it greatly overlaps with beta C1 found in HC, with a decrease in connectivity before RT (although stopping much earlier) and an increase beginning around averaged RT and that was sustained after. Therefore, at approximately the same timing, functional brain states were spatially different at the group-level between PD and HC. In addition, a second dBNS was found in PD patients with a significant increase in connectivity that began before, and that overlapped with, the averaged RT; and which was absent in HC. This dBNS was mostly left-sided with a strong temporal component, but which also included CAC-associated regions such as inferior frontal and orbitofrontal regions as well as the anterior cingulate. One could wonder if the strong lateralization of those networks is linked to the number of left- or right-presented stimuli. In that respect, we checked and found a significant difference with more left than right stimuli trials that remained for EEG analyses. However the effect size was small, and as a consequence we considered it as negligible (Cohen’s d = 0.2). It is thus unlikely that the stimulus’ side fully explains the lateralization of the networks that we identified.

One surprising aspect is that we would expect PD-related beta changes to involve motor regions in PD patients. PD has been indeed consistently associated with increased beta power in motor circuits (Little and Brown, 2014). One explanation of the absence of motor nodes might lie in the fact that we used phase-based functional connectivity. Oscillatory phase is, to some extent, independent from amplitude. Thus, local increase in beta power wouldn’t necessarily be associated with phase synchronization between motor and other areas. Following this, it is also possible that such beta phase synchronization actually occurred, but was arguably too short-lived to be highlighted by temporal ICA.

### 4.3. Several gamma networks overlap in PD

Regarding the gamma band, we found two significant and spatially distinct dBNS in HC. The first one occurred before the average RT and involved CAC-related left frontal areas such as the inferior frontal, anterior cingulate, and orbitofrontal regions. These regions connected right temporal ones. The second dBNS occured after the average RT, and also involved the same CAC-related frontal region, albeit in a more bilateral way than observed in gamma C1. In this second dBNS, the temporal component was also present, but on the left and to a lesser extent. In PD patients, the 3 dBNS that we found were spatially different from HC and had a more complex temporal organization. One dBNS showed increased connectivity before, and overlapping with, the average RT. It included CAC-related frontal areas, including inferior frontal, anterior cingulate and orbitofrontal regions; and also showed temporo-temporal connections. The two other dBNS were overlapping in time with the first dBNS and with themselves, and had decreased connectivity around the average RT with long-range fronto-temporal and fronto-occipital connections. In HC, no such fronto-occipital connectivity was found. Overall, although sharing some CAC-related prefrontal nodes (also described in beta) with HC, the PD networks were spatially different. It is interesting to note that the prefrontal component that included most CAC-related regions in gamma C2 was very similar to the one in beta C1.

Similarly to beta oscillations, gamma oscillations have been associated with several cognitive functions (see (Herrmann et al., 2010) for a review). Unfortunately, studies reporting gamma effects on cognitive action control are scarce. However, some found increased gamma activity in a multi-source interference task in somato-sensory and occipital regions (Wiesman et al., 2020; Wiesman and Wilson, 2020). Among the various roles attributed to gamma oscillations, binding has probably been the most discussed. The binding theory hypothesizes that different stimulus features (color, spatial location, shape, etc.) are ultimately binded into a single representation by the means of gamma synchronization (Engel et al., 2001; Singer, 1999). Synchronization would then be necessary for attention, sensori-motor integration and response selection. In that respect, long-range gamma connectivity, as we observed in our various brain network states, would reflect the binding of sensory and cognitive information used to perform the task. Assigning a cognitive role to gamma oscillations is nonetheless debated, and some researchers argue that gamma activity (and long-range gamma synchrony) do not necessarily have a functional role in cognition, but rather reflect local states of activation (Merker, 2016; Ray and Maunsell, 2015). Regarding PD effect on gamma oscillations, motor and frontal gamma typically show coherent activity with subcortical structures (such as the subthalamic nucleus) and might modulate the vigor of a motor response (Oswal et al., 2013). However, the gamma activity usually described in PD studies spans higher and larger gamma bands (60-90 Hz) than the one we used here (30-45 Hz). Also, although we found prefrontal gamma in our brain network states, no motor nodes were present.

### 4.4. Network presence differed between PD and HC at the subject-level

The use of typical microstate metrics allowed us to investigate the presence of all dBNS at the subject-level. Several metrics were used that inform on the lifespan, frequency of occurrence, coverage time, and explained variance of dBNS, as well as transition probability between states. An important point to bear in mind is that the computation of these metrics is based on the similarity between the dFC matrices of a subject, and the dBNS. At each time, one dBNS is deemed the most similar to the dFC matrix based on a spatial correlation. Consequently, for all dFC matrices, only one dBNS is set to be the most similar, although other dBNS can be present. This explains why some dBNS significantly present at the group-level had 0 values at the subject-level for the network states metrics.

Differences between PD patients and HC were mostly found in the beta band. Globally, the C1 dBNS dominated in both groups (although originally defined only in HC) but was more present in HC. Importantly, the frequency of occurrence of C1 was associated with congruent and incongruent RT in such a way that increased presence of C1 was linked to decreased RT. This result, although purely correlational, is interesting because C1 was less present in PD and patients had longer RT. The P2 dBNS was the second dominant brain state, while P1 never dominated, and was more present and explained more variance in PD patients. In the gamma band, the difference between PD patients and HC was weaker and only supported by a significant decrease in the frequency of occurrence of P2 in PD patients as compared to HC. It is important to note that this network, although originally defined on PD data, dominated in both groups at the subject-level and that HC even showed an increased frequency of occurrence on that dBNS.

### 4.5. Methodological considerations and limitations

First, we focused our EEG analyses on correct incongruent trials only. At the behavioral level, CAC effects are traditionally obtained by contrasting the congruent and incongruent situations. Regarding dynamic functional brain states, it is not possible to simply subtract signals between congruent and incongruent trials to obtain networks that reflect this behavioral difference. We chose to focus on correct incongruent trials, since they reflect the implementation of CAC in the situation of highest conflict during the task. Our interpretations are thus limited to CAC in the incongruent situation, and do not reflect the traditional congruence effect observed in the behavioral data.

Our estimation of dynamic brain network states depended on several parameters that can be discussed, and must be kept in mind for interpreting the results. For instance, determining the optimal number of derived components is still a challenging question for most decomposition algorithms, including tICA. In this study, the selection of the number of states was performed in two consecutive steps: (1) a primary number of components was selected using the difference in data fitting (DIFFIT) technique adopted by previous studies (Timmerman and Kiers, 2000; Wang et al., 2018). Then, (2) given that all brain states are not necessarily associated with the task, we searched for components that were significantly modulated by the task. We used permutation testing by building a non-parametric null distribution generated using a ‘sign-flipping’ approach (Hunt et al., 2012; O’Neill et al., 2017; Winkler et al., 2014). This approach highlighted the components with trial-averaged time courses that exceeded a specific threshold. The use of the null distribution may be deemed as too conservative, since a Bonferroni correction for multiple comparisons was carried out, as well as a constraint on the minimal number of significant cycles (3). For example, we obtained only one significant component for HC and two for PD in the beta band. It could be argued that such a restrictive number of components might miss additional information. However, we did not have any a priori hypothesis on the number of brain states in PD or HC, nor on their duration. Therefore, a conservative approach appears more appropriate.

We used a backfitting approach to estimate typical microstate metrics for subject-level analyses. Although this method is practical to estimate the presence of brain states at the subject-level, it suffers from a drawback which consists in assuming that only one global functional state occurs at a given moment in time. Thus, estimation of these metrics is based on the determination of one dominant dBNS at each time. This completely ignores the assumption of temporal ICA, which considers overlapping and independent brain network states with proportional weights to summarize brain activity. As a consequence, our interpretations of subject-level analyses are limited by the fact that we focused on dominant dBNS. Future studies could explore further other back-fitting approaches that rely on the notion of proportional, rather than binary, fitting.

To estimate the spatial similarity between the dBNS and dFC matrices, we used a simple correlation measure at the back-fitting step. We chose this measure since we aimed to assign to each functional network the nearest (spatially) dBNS, rather than evaluating the precision of the exact similarity value by itself. Nevertheless, other spatial similarity metrics could be used in this context, such as those taking into account the spatial locations of the compared networks nodes (Mheich et al., 2018; Pineda-Pardo et al., 2015), or distance-based metrics (Cao et al., 2013; Gao et al., 2010).

Another important point, regarding our subject-level results, one could argue that it is not surprising that dBNS derived from PD (as beta P2) are more present at the subject-level in PD patients than in HC. This is indeed a limitation to bear in mind while interpreting our subject-level results. We chose to apply tICA in a specific manner for the PD and HC group, because we aimed to highlight group-level spatial and temporal differences between PD patients and HC. Consequently, it is not surprising that the correlation between a group-specific dBNS correlates with a subject’s dFC matrices in that same group. However, this limitation is lessened by the fact that some dBNS defined in PD, such as gamma P2, were also dominant in HC at the subject-level. We even observed a significantly higher frequency of occurrence in HC for gamma P2, although it was defined in the PD group.

Finally, it is worth noting that investigation of dynamic functional connectivity using the methods reported here drastically constrain the analyses to relatively high frequencies. Indeed, performing the same analyses in frequencies lower than beta would have been challenging, given the limited number of cycles available in the duration of the trial. This is a key point, since most CAC-related results in the literature involve theta frequencies and report increased midfrontal theta activity during conflict (Cohen, 2014). One interesting perspective would be to investigate if the dynamic presence of certain beta brain states in PD depend on fronto-central theta activity, since midfrontal theta-parietal beta cross-frequency coupling has been shown during CAC (Duprez et al., 2020) and PD appears to be associated with decreased midfrontal theta (Singh et al., 2018).

## 5. Conclusion

In this study, we reported brain functional connectivity states at the source level that differed between PD patients and healthy controls during conflict situations. We showed, both at the group- and subject-level, that a strong prefrontal beta component found in HC was less present in PD, which correlated with reaction time at the task. We also highlighted that several brain networks in the gamma band overlapped in PD. Although the method used allowed for an estimation of dominant dynamic brain states, it might not be appropriate in the evaluation of subject-level overlapping networks. Nevertheless, we believe that this study highlights the relevance of task-based dynamic connectivity measures which could help in the understanding of cognitive dysfunctions observed in PD, and in other neurological diseases.

## Authorship contribution statement

JD: Conceptualization, Methodology, Software, Data curation, Writing – original draft, Visualization, Investigation, Validation, Writing – review & editing. JT: Methodology, Software, Data curation, Writing – original draft, Visualization, Writing – review & editing. MH: Conceptualization, Methodology, Writing – review & editing. JM: Methodology, Writing – review & editing. AK: Writing – review & editing. AM: Writing – review & editing. SD: Writing – review & editing. MV: Writing – review & editing. PS: Conceptualization, Writing – review & editing. FW: Writing – review & editing. PB: Conceptualization, Methodology, Data curation, Investigation, Validation, Writing – review & editing. JFH: Conceptualization, Methodology, Data curation, Investigation, Validation, Writing – review & editing.

## Acknowledgements

The authors would like to thank all the participants who took part in this study. We also thank Dr. Adrien Bénard (MD) and Dr. Sina Potel (MD) for their help in data acquisition. We also would like to thank Bretagne Atlantique Ambition (BAA) as well as the Rennes Clinical Neuroscience Institute (INCR: www.incr.fr) who funded this work.

## Funding

JD was funded by the Rennes Clinical Neuroscience Institute (INCR: www.incr.fr).

## Declarations of interest

None.

## Abbreviations

CAC: Cognitive action control
dBNS: Dynamic brain network state
dFC: Dynamic functional connectivity
DIFFIT: Difference in data fitting
DLPFC: Dorso-lateral prefrontal cortex
EEG: Electroencephalography
FC: Functional connectivity
fMRI: Functional magnetic resonance imaging
HC: Healthy controls
HD-EEG: High-density
EEG ICA: Independent component analysis
IFC: Inferior frontal cortex
MEG: Magnetoencephalography
PD: Parkinson’s disease
PLV: Phase locking value
pre-SMA: Pre-supplementary motor area
ROIS: Regions of interest
RT: Reaction time
tICA: Temporal ICA

## Supplementary Materials

**Supplementary Table 1 :**
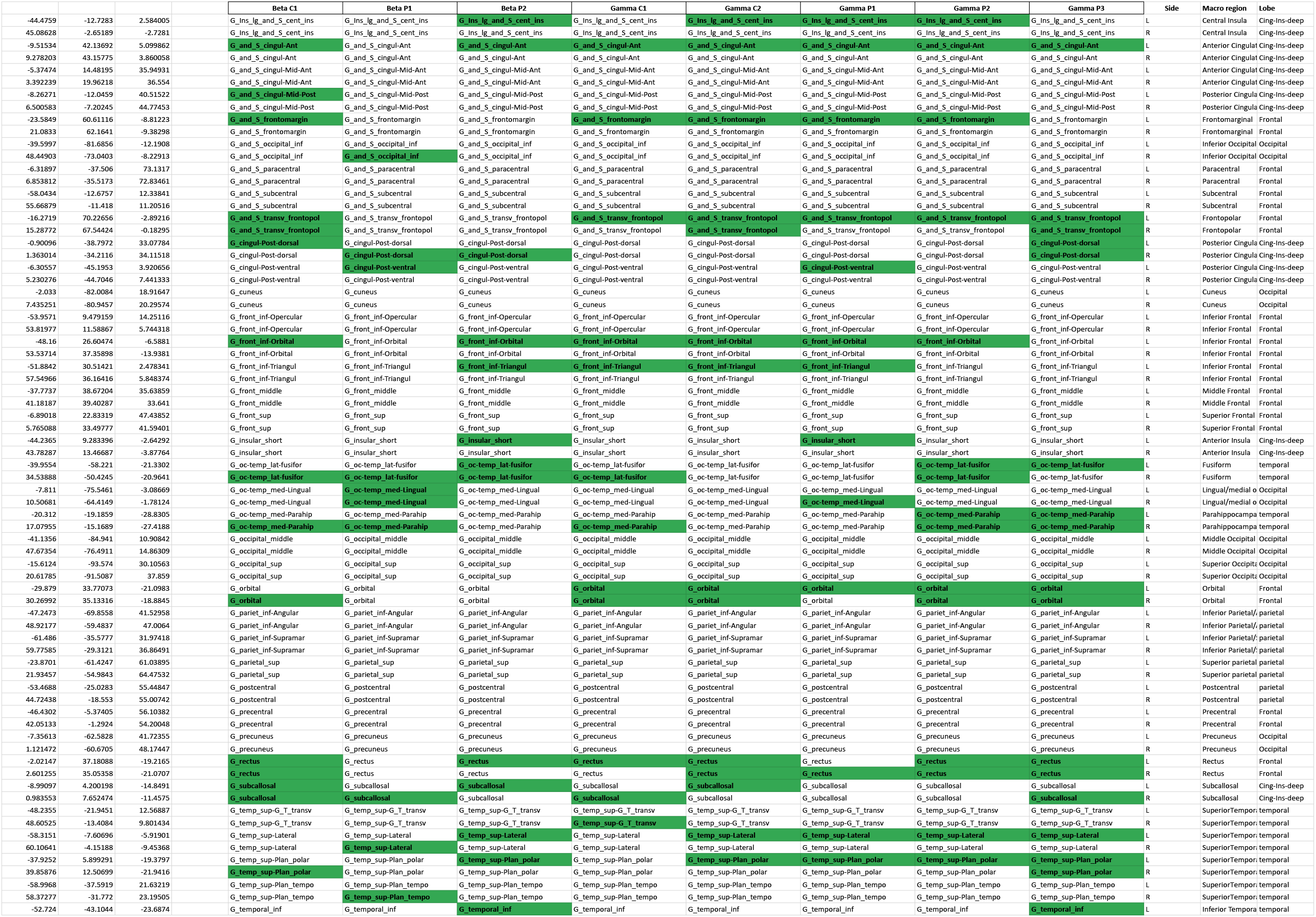
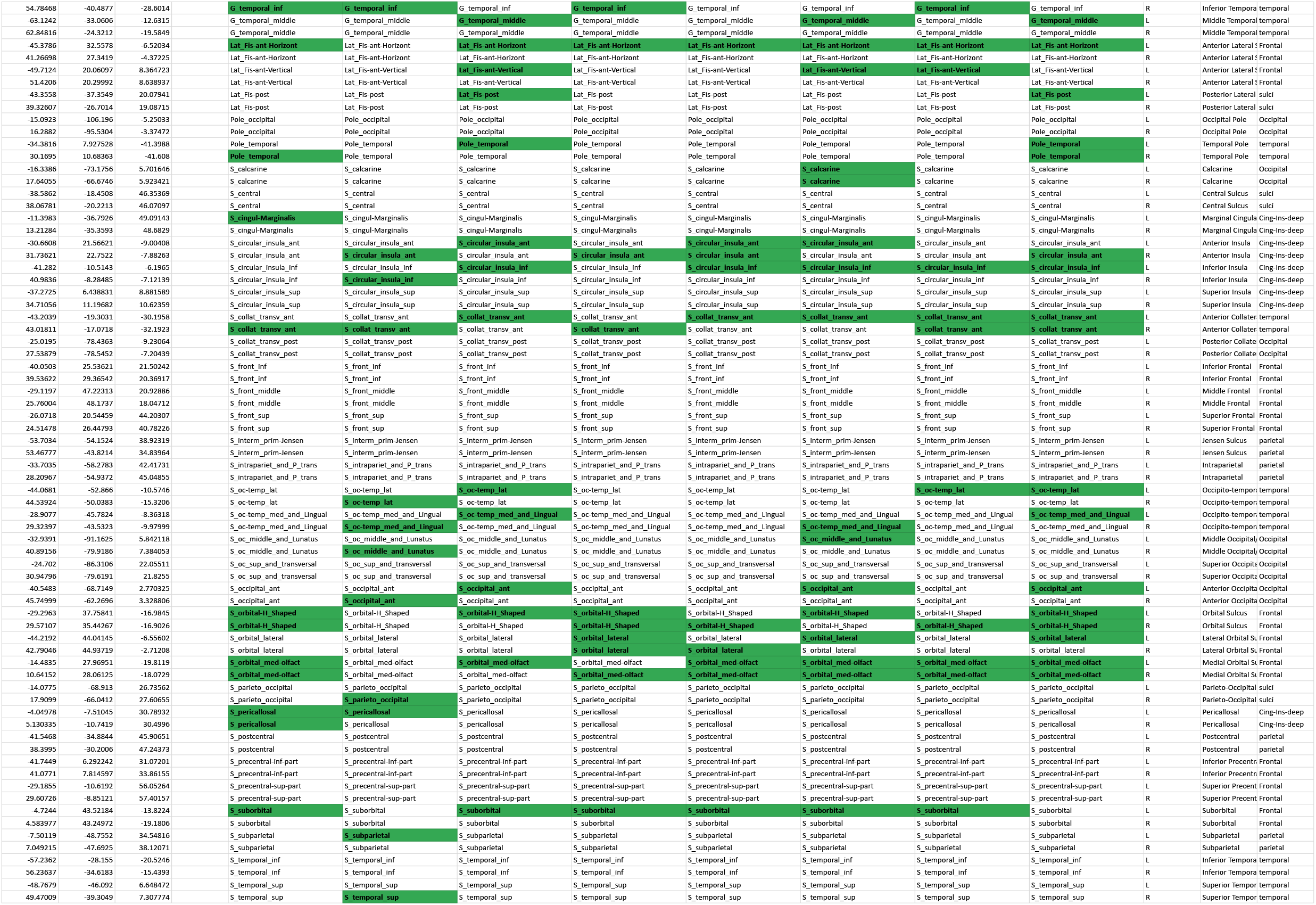
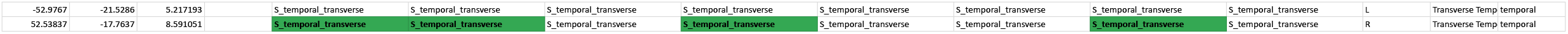
Destrieux atlas with all nodes found in the HC and PD dBNS. Nodes present in the networks are highlighted in green.

**Supplementary Figure 1.**
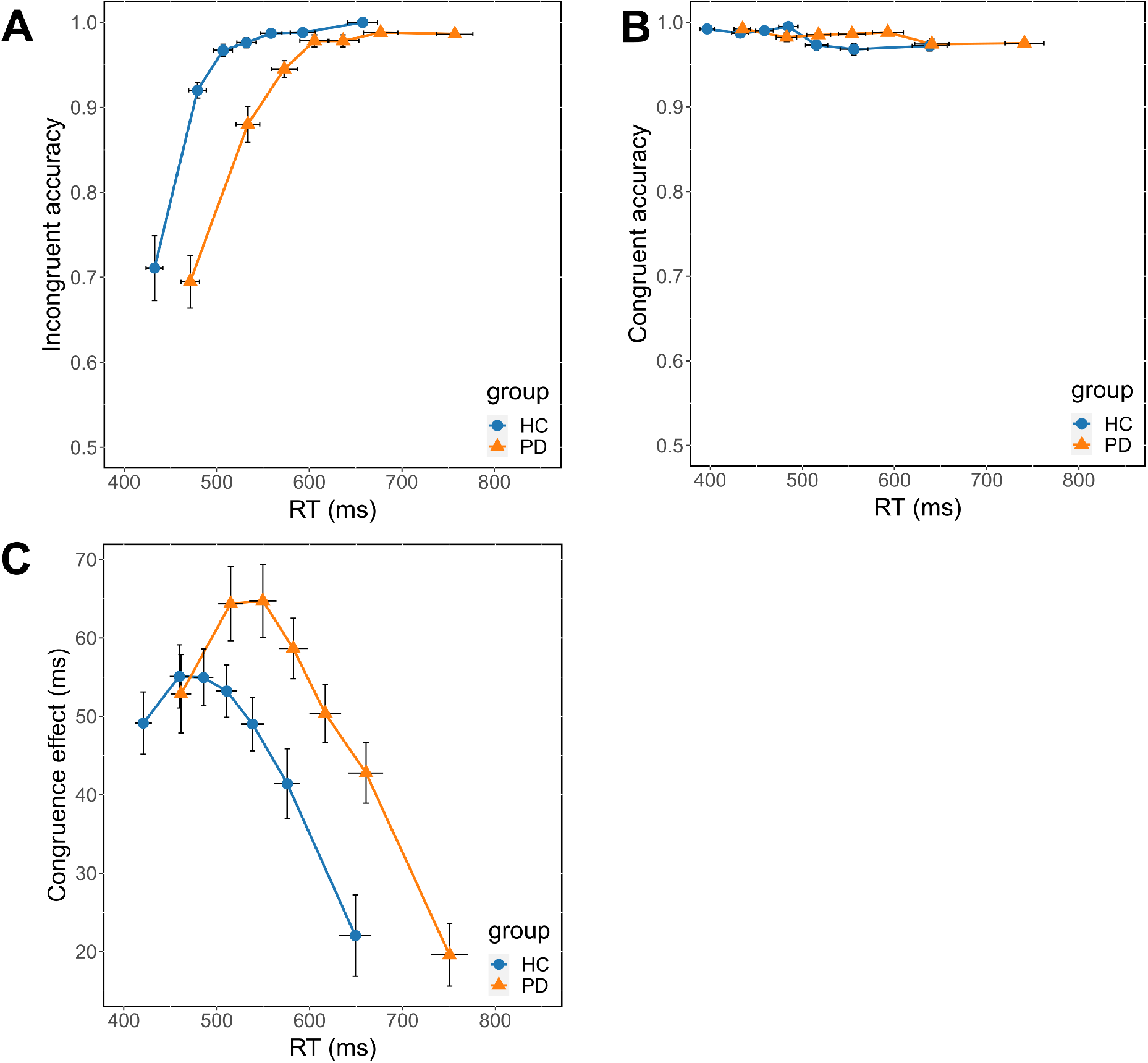
Conditional accuracy functions representing accuracy as a function of the RT distribution in the incongruent (A) and congruent (B) situations. C: Delta plots showing the evolution of the congruence effect as a function of the RT distribution.

**Supplementary Figure 2.**
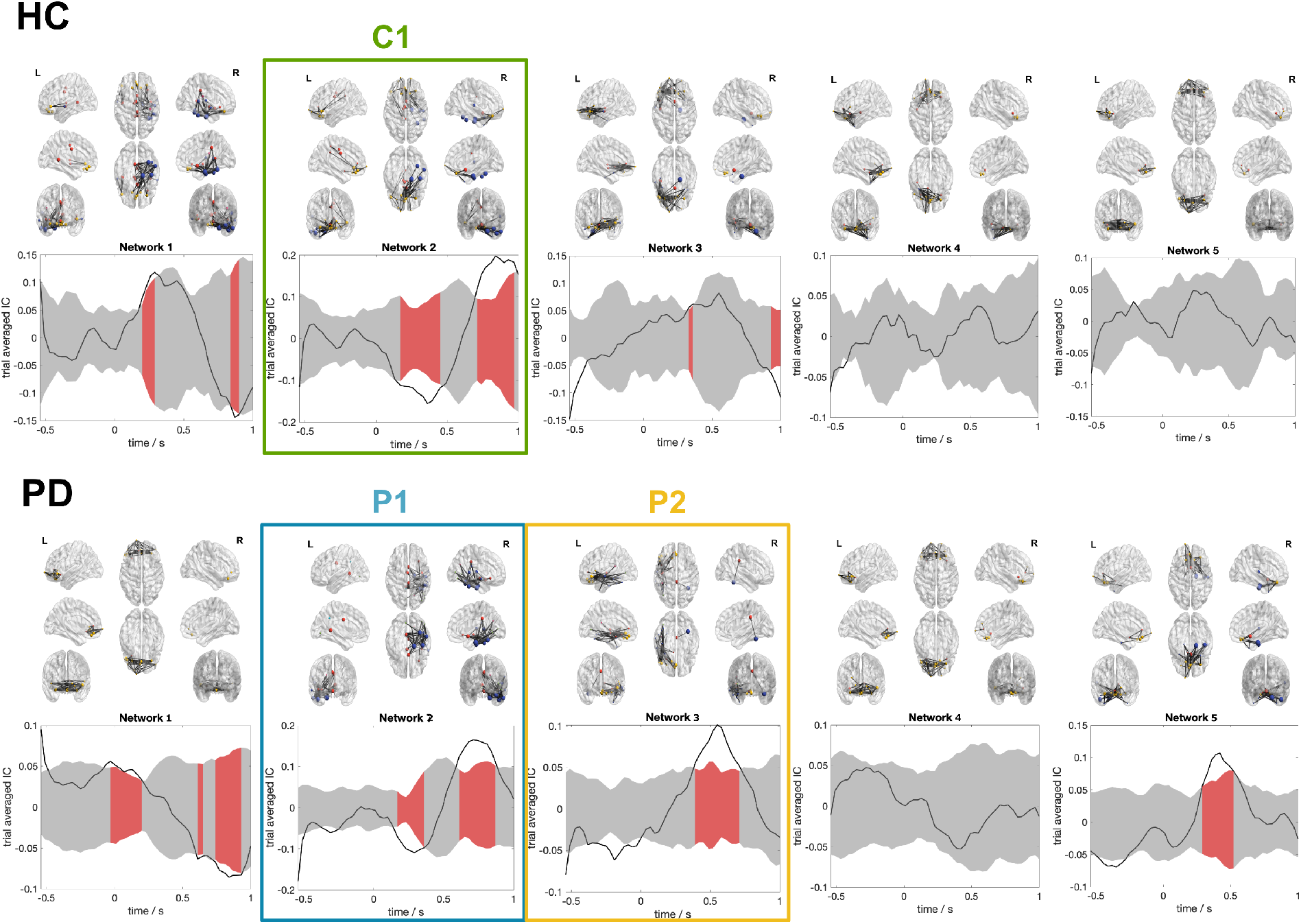
All dBNS extracted via tICA in HC and PD in the beta band. Cortical networks 3D representations are followed by the evolution of presence during the time of a trial. Null distributions are plotted in gray-shaded areas. Periods of significance (above or below the null distribution) are shaded in red.

**Supplementary Figure 3.**
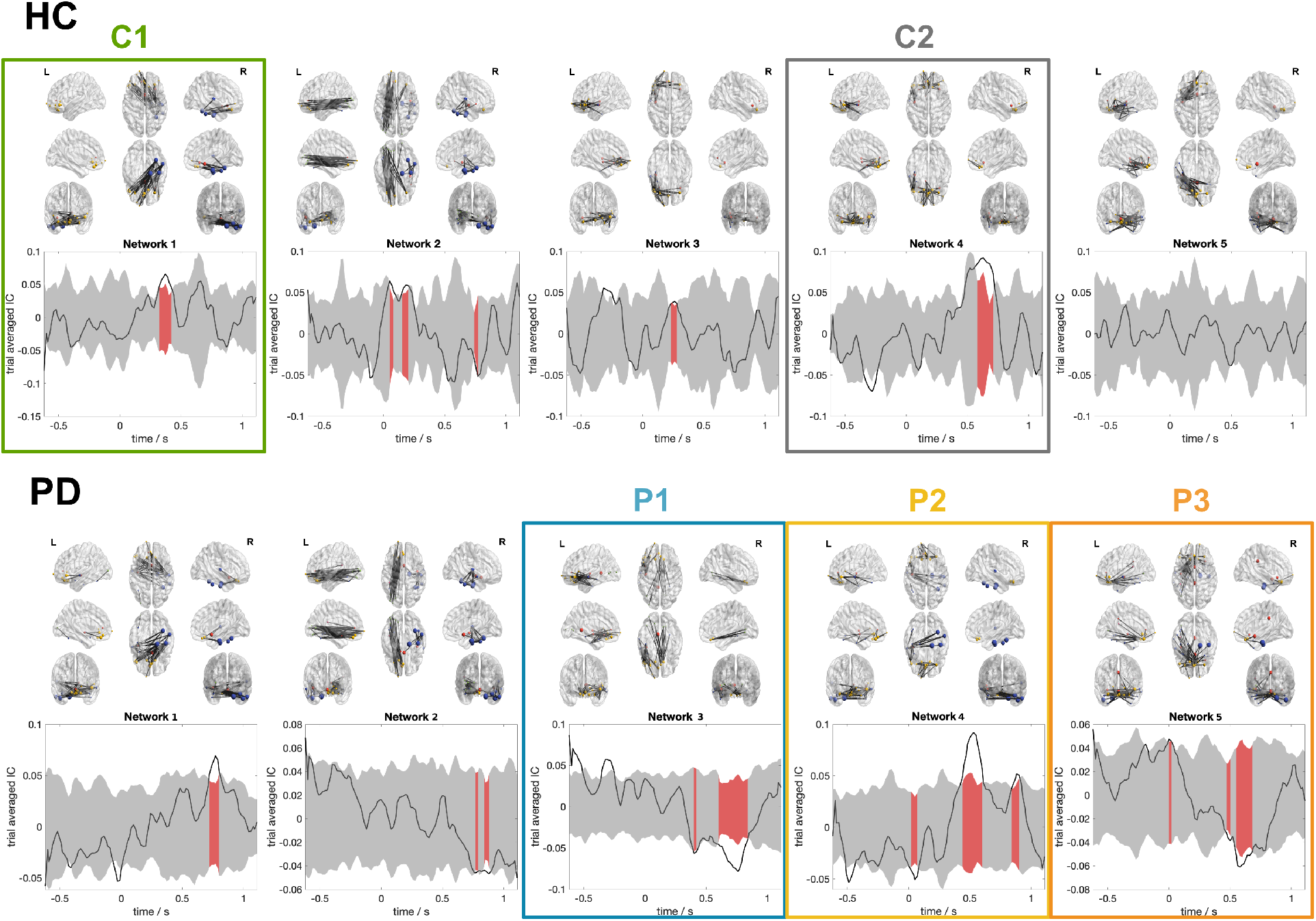
All dBNS extracted via tICA in HC and PD in the gamma band. Cortical networks 3D representations are followed by the evolution of presence during the time of a trial. Null distributions are plotted in gray-shaded areas. Periods of significance (above or below the null distribution) are shaded in red.

**Supplementary Figure 4.**
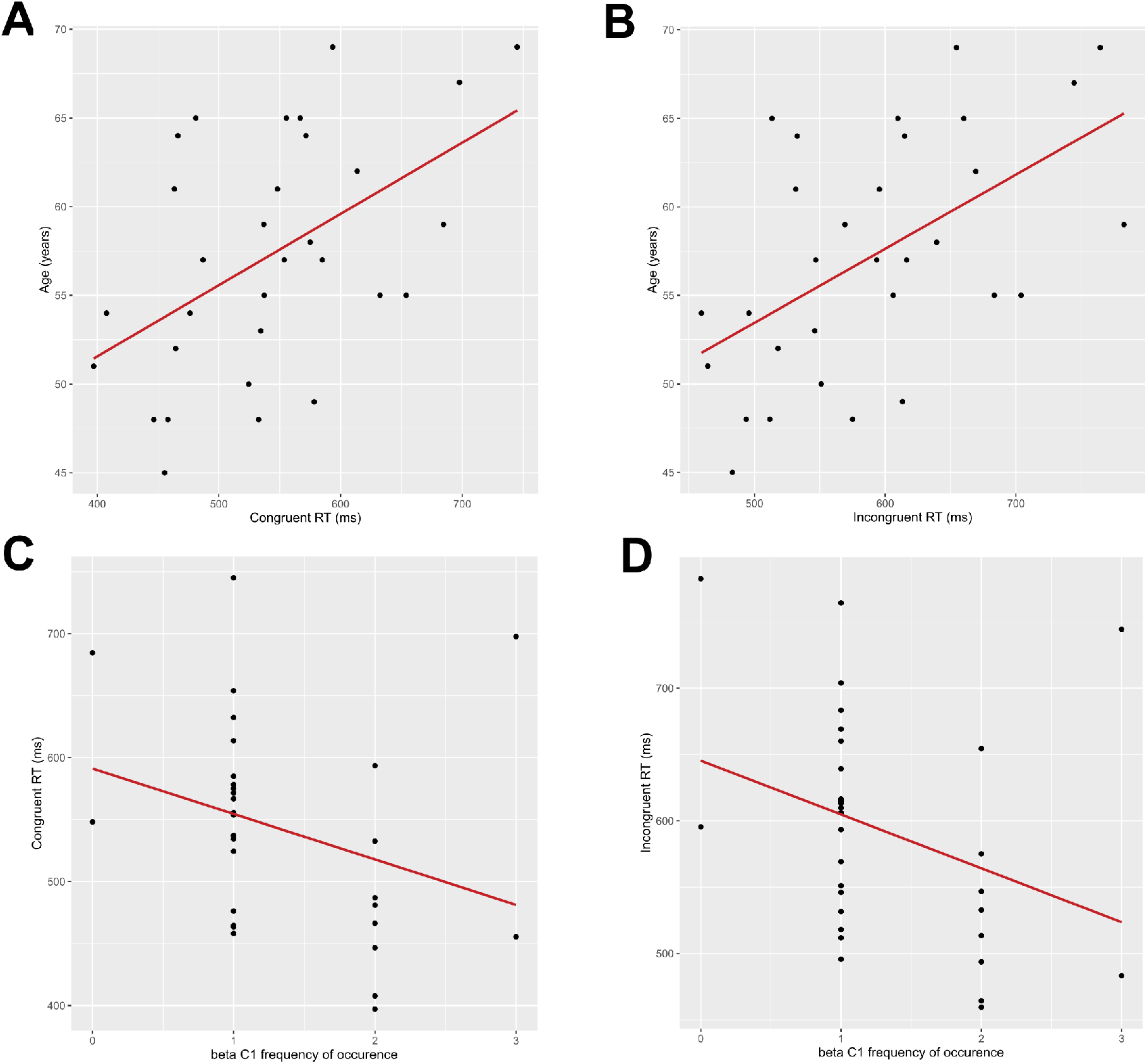
Scatter plots showing the correlation between age and congruent (A) and incongruent (B) RT. C and D show the relations between the frequency of occurence of the beta C1 dBNS and congruent and incongruent RT respectively.

## Notes

https://github.com/judytabbal/dynCogPD

### Competing Interest Statement

The authors have declared no competing interest.

## References

Allen, E.A., Damaraju, E., Eichele, T., Wu, L., Calhoun, V.D., 2018. EEG Signatures of Dynamic Functional Network Connectivity States. Brain Topogr. 31, 101–116. https://doi.org/10.1007/s10548-017-0546-2

Aron, A.R., Robbins, T.W., Poldrack, R.A., 2004. Inhibition and the right inferior frontal cortex. Trends Cogn. Sci. 8, 170–177. https://doi.org/10.1016/j.tics.2004.02.010

Baggio, H.-C., Sala-Llonch, R., Segura, B., Marti, M.-J., Valldeoriola, F., Compta, Y., Tolosa, E., Junqué, C., 2014. Functional brain networks and cognitive deficits in Parkinson’s disease. Hum. Brain Mapp. 35, 4620–4634. https://doi.org/10.1002/hbm.22499

Baggio, H.-C., Segura, B., Sala-Llonch, R., Marti, M.-J., Valldeoriola, F., Compta, Y., Tolosa, E., Junqué, C., 2015. Cognitive impairment and resting-state network connectivity in Parkinson’s disease. Hum. Brain Mapp. 36, 199–212. https://doi.org/10.1002/hbm.22622

Baker, A.P., Brookes, M.J., Rezek, I.A., Smith, S.M., Behrens, T., Probert Smith, P.J., Woolrich, M., 2014. Fast transient networks in spontaneous human brain activity. eLife 3, e01867. https://doi.org/10.7554/eLife.01867

Barton, K., 2009. MuMIn: multi-model inference. Httpr-Forge R-Proj. Orgprojectsmumin.

Bassett, D.S., Sporns, O., 2017. Network neuroscience. Nat. Neurosci. 20, 353–364. https://doi.org/10.1038/nn.4502

Bates, D., Mächler, M., Bolker, B., Walker, S., 2015. Fitting Linear Mixed-Effects Models Using lme4. J. Stat. Softw. 67, 1–48. https://doi.org/10.18637/jss.v067.i01

Benton, A.L., Varney, N.R., Hamsher, K. deS, 1978. Visuospatial judgment: A clinical test. Arch. Neurol. 35, 364–367.

Bola, M., Sabel, B.A., 2015. Dynamic reorganization of brain functional networks during cognition. NeuroImage 114, 398–413. https://doi.org/10.1016/j.neuroimage.2015.03.057

Bridi, J.C., Hirth, F., 2018. Mechanisms of -Synuclein Induced Synaptopathy in Parkinson’s Disease. Front. Neurosci. 12, 80. https://doi.org/10.3389/fnins.2018.00080

Brodbeck, V., Kuhn, A., von Wegner, F., Morzelewski, A., Tagliazucchi, E., Borisov, S., Michel, C.M., Laufs, H., 2012. EEG microstates of wakefulness and NREM sleep. NeuroImage 62, 2129–2139. https://doi.org/10.1016/j.neuroimage.2012.05.060

Cagigas, X.E., Filoteo, J.V., Stricker, J.L., Rilling, L.M., Friedrich, F.J., 2007. Flanker compatibility effects in patients with Parkinson’s disease: impact of target onset delay and trial-by-trial stimulus variation. Brain Cogn. 63, 247–259. https://doi.org/10.1016/j.bandc.2006.09.002

Cao, B., Li, Y., Yin, J., 2013. Measuring Similarity between Graphs Based on the Levenshtein Distance. Appl. Math. Inf. Sci. 7, 169–175. https://doi.org/10.12785/amis/071L24

Cardoso, J.F., Souloumiac, A., 1993. An efficient technique for the blind separation of complex sources, in: [1993 Proceedings] IEEE Signal Processing Workshop on Higher-Order Statistics. Presented at the [1993 Proceedings] IEEE Signal Processing Workshop on Higher-Order Statistics, pp. 275–279. https://doi.org/10.1109/HOST.1993.264552

Cespón, J., Hommel, B., Korsch, M., Galashan, D., 2020. The neurocognitive underpinnings of the Simon effect: An integrative review of current research. Cogn. Affect. Behav. Neurosci. 20, 1133–1172. https://doi.org/10.3758/s13415-020-00836-y

Cohen, M.X., 2014. A neural microcircuit for cognitive conflict detection and signaling. Trends Neurosci. 37, 480–490. https://doi.org/10.1016/j.tins.2014.06.004

Cong, F., He, Z., Hämäläinen, J., Leppänen, P.H.T., Lyytinen, H., Cichocki, A., Ristaniemi, T., 2013. Validating rationale of group-level component analysis based on estimating number of sources in EEG through model order selection. J. Neurosci. Methods 212, 165–172. https://doi.org/10.1016/j.jneumeth.2012.09.029

Dale, A.M., Sereno, M.I., 1993. Improved Localizadon of Cortical Activity by Combining EEG and MEG with MRI Cortical Surface Reconstruction: A Linear Approach. J. Cogn. Neurosci. 5, 162–176. https://doi.org/10.1162/jocn.1993.5.2.162

de Pasquale, F., Della Penna, S., Sporns, O., Romani, G.L., Corbetta, M., 2016. A Dynamic Core Network and Global Efficiency in the Resting Human Brain. Cereb. Cortex 26, 4015–4033. https://doi.org/10.1093/cercor/bhv185

Destrieux, C., Fischl, B., Dale, A., Halgren, E., 2010. Automatic parcellation of human cortical gyri and sulci using standard anatomical nomenclature. NeuroImage 53, 1–15. https://doi.org/10.1016/j.neuroimage.2010.06.010

Duprez, J., Gulbinaite, R., Cohen, M.X., 2020. Midfrontal theta phase coordinates behaviorally relevant brain computations during cognitive control. NeuroImage 207, 116340. https://doi.org/10.1016/j.neuroimage.2019.116340

Duprez, J., Houvenaghel, J.-F., Argaud, S., Naudet, F., Robert, G., Drapier, D., Vérin, M., Sauleau, P., 2017. Impulsive oculomotor action selection in Parkinson’s disease. Neuropsychologia 95, 250–258. https://doi.org/10.1016/j.neuropsychologia.2016.12.027

Eagle, D.M., Baunez, C., Hutcheson, D.M., Lehmann, O., Shah, A.P., Robbins, T.W., 2008. Stop-Signal Reaction-Time Task Performance: Role of Prefrontal Cortex and Subthalamic Nucleus. Cereb. Cortex 18, 178–188. https://doi.org/10.1093/cercor/bhm044

Engel, A.K., Fries, P., 2010. Beta-band oscillations–signalling the status quo? Curr. Opin. Neurobiol. 20, 156–165. https://doi.org/10.1016/j.conb.2010.02.015

Engel, A.K., Fries, P., Singer, W., 2001. Dynamic predictions: Oscillations and synchrony in top–down processing. Nat. Rev. Neurosci. 2, 704–716. https://doi.org/10.1038/35094565

Falkenstein, M., Willemssen, R., Hohnsbein, J., Hielscher, H., 2006. Effects of stimulus-response compatibility in Parkinson’s disease: a psychophysiological analysis. J. Neural Transm. 113, 1449–1462.

Fornito, A., Zalesky, A., Breakspear, M., 2015. The connectomics of brain disorders. Nat. Rev. Neurosci. 16, 159–172. https://doi.org/10.1038/nrn3901

Forstmann, B., van den Wildenberg, W., Ridderinkhof, K., 2008. Neural mechanisms, temporal dynamics, and individual differences in interference control. Cogn. Neurosci. J. Of 20, 1854–1865.

Forstmann, B.U., Jahfari, S., Scholte, H.S., Wolfensteller, U., van den Wildenberg, W.P., Ridderinkhof, K.R., 2008. Function and structure of the right inferior frontal cortex predict individual differences in response inhibition: a model-based approach. J. Neurosci. 28, 9790–9796.

Fox, J., Weisberg, S., 2019. An R Companion to Applied Regression, Third. ed. Sage, Thousand Oaks CA.

Friedman, N.P., Robbins, T.W., 2021. The role of prefrontal cortex in cognitive control and executive function. Neuropsychopharmacology 1–18. https://doi.org/10.1038/s41386-021-01132-0

Fries, P., 2015. Rhythms for cognition: communication through coherence. Neuron 88, 220–235.

Gao, X., Xiao, B., Tao, D., Li, X., 2010. A survey of graph edit distance. Pattern Anal. Appl. 13, 113–129. https://doi.org/10.1007/s10044-008-0141-y

Gramfort, A., Papadopoulo, T., Olivi, E., Clerc, M., 2010. OpenMEEG: opensource software for quasistatic bioelectromagnetics. Biomed. Eng. OnLine 9, 45. https://doi.org/10.1186/1475-925X-9-45

Graves, R.E., Bezeau, S.C., Fogarty, J., Blair, R., 2004. Boston naming test short forms: a comparison of previous forms with new item response theory based forms. J. Clin. Exp. Neuropsychol. 26, 891–902.

Haber, S.N., 2014. The place of dopamine in the cortico-basal ganglia circuit. Neuroscience, The Ventral Tegmentum and Dopamine: A New Wave of Diversity 282, 248–257. https://doi.org/10.1016/j.neuroscience.2014.10.008

Hämäläinen, M.S., Ilmoniemi, R.J., 1994. Interpreting magnetic fields of the brain: minimum norm estimates. Med. Biol. Eng. Comput. 32, 35–42. https://doi.org/10.1007/BF02512476

Hassan, M., Benquet, P., Biraben, A., Berrou, C., Dufor, O., Wendling, F., 2015. Dynamic reorganization of functional brain networks during picture naming. Cortex J. Devoted Study Nerv. Syst. Behav. 73, 276–288. https://doi.org/10.1016/j.cortex.2015.08.019

Hassan, M., Wendling, F., 2018. Electroencephalography Source Connectivity: Aiming for High Resolution of Brain Networks in Time and Space. IEEE Signal Process. Mag. 35, 81–96. https://doi.org/10.1109/MSP.2017.2777518

Hayes, M.T., 2019. Parkinson’s Disease and Parkinsonism. Am. J. Med. 132, 802–807. https://doi.org/10.1016/j.amjmed.2019.03.001

Herrmann, C.S., Fründ, I., Lenz, D., 2010. Human gamma-band activity: a review on cognitive and behavioral correlates and network models. Neurosci. Biobehav. Rev. 34, 981–992. https://doi.org/10.1016/j.neubiorev.2009.09.001

Hommel, B., Wiers, R.W., 2017. Towards a Unitary Approach to Human Action Control. Trends Cogn. Sci. 21, 940–949. https://doi.org/10.1016/j.tics.2017.09.009

Hughes, A.J., Ben-Shlomo, Y., Daniel, S.E., Lees, A.J., 1992. What features improve the accuracy of clinical diagnosis in Parkinson’s disease: a clinicopathologic study. Neurology 42, 1142–1146.

Hunt, L.T., Kolling, N., Soltani, A., Woolrich, M.W., Rushworth, M.F.S., Behrens, T.E.J., 2012. Mechanisms underlying cortical activity during value-guided choice. Nat. Neurosci. 15, 470–476. https://doi.org/10.1038/nn.3017

Kabbara, A., Paban, V., Hassan, M., 2021. The dynamic modular fingerprints of the human brain at rest. NeuroImage 227, 117674. https://doi.org/10.1016/j.neuroimage.2020.117674

Khanna, A., Pascual-Leone, A., Michel, C.M., Farzan, F., 2015. Microstates in resting-state EEG: Current status and future directions. Neurosci. Biobehav. Rev. 49, 105–113. https://doi.org/10.1016/j.neubiorev.2014.12.010

Kim, C., Johnson, N.F., Gold, B.T., 2012. Common and distinct neural mechanisms of attentional switching and response conflict. Brain Res. 1469, 92–102. https://doi.org/10.1016/j.brainres.2012.06.013

Knyazev, G.G., Savostyanov, A.N., Bocharov, A.V., Tamozhnikov, S.S., Saprigyn, A.E., 2016. Task-positive and task-negative networks and their relation to depression: EEG beamformer analysis. Behav. Brain Res. 306, 160–169. https://doi.org/10.1016/j.bbr.2016.03.033

Koelewijn, L., Bompas, A., Tales, A., Brookes, M.J., Muthukumaraswamy, S.D., Bayer, A., Singh, K.D., 2017. Alzheimer’s disease disrupts alpha and beta-band resting-state oscillatory network connectivity. Clin. Neurophysiol. 128, 2347–2357. https://doi.org/10.1016/j.clinph.2017.04.018

Koelewijn, L., Hamandi, K., Brindley, L.M., Brookes, M.J., Routley, B.C., Muthukumaraswamy, S.D., Williams, N., Thomas, M.A., Kirby, A., te Water Naudé, J., Gibbon, F., Singh, K.D., 2015. Resting-state oscillatory dynamics in sensorimotor cortex in benign epilepsy with centro-temporal spikes and typical brain development. Hum. Brain Mapp. 36, 3935–3949. https://doi.org/10.1002/hbm.22888

Kötter, R., Mazziotta, J., Toga, A., Evans, A., Fox, P., Lancaster, J., Zilles, K., Woods, R., Paus, T., Simpson, G., Pike, B., Holmes, C., Collins, L., Thompson, P., MacDonald, D., Iacoboni, M., Schormann, T., Amunts, K., Palomero-Gallagher, N., Geyer, S., Parsons, L., Narr, K., Kabani, N., Goualher, G.L., Boomsma, D., Cannon, T., Kawashima, R., Mazoyer, B., 2001. A probabilistic atlas and reference system for the human brain: International Consortium for Brain Mapping (ICBM). Philos. Trans. R. Soc. Lond. B. Biol. Sci. 356, 1293–1322. https://doi.org/10.1098/rstb.2001.0915

Kulashekhar, S., Pekkola, J., Palva, J.M., Palva, S., 2016. The role of cortical beta oscillations in time estimation. Hum. Brain Mapp. 37, 3262–3281. https://doi.org/10.1002/hbm.23239

Lachaux, J.-P., Rodriguez, E., Le Van Quyen, M., Lutz, A., Martinerie, J., Varela, F.J., 2000. Studying single-trials of phase synchronous activity in the brain. Int. J. Bifurc. Chaos 10, 2429–2439. https://doi.org/10.1142/S0218127400001560

Lawson, R.A., Yarnall, A.J., Duncan, G.W., Breen, D.P., Khoo, T.K., Williams-Gray, C.H., Barker, R.A., Collerton, D., Taylor, J.-P., Burn, D.J., ICICLE-PD study group, 2016. Cognitive decline and quality of life in incident Parkinson’s disease: The role of attention. Parkinsonism Relat. Disord. 27, 47–53. https://doi.org/10.1016/j.parkreldis.2016.04.009

Lehmann, D., Faber, P.L., Galderisi, S., Herrmann, W.M., Kinoshita, T., Koukkou, M., Mucci, A., Pascual-Marqui, R.D., Saito, N., Wackermann, J., Winterer, G., Koenig, T., 2005. EEG microstate duration and syntax in acute, medication-naïve, first-episode schizophrenia: a multi-center study. Psychiatry Res. Neuroimaging 138, 141–156. https://doi.org/10.1016/j.pscychresns.2004.05.007

Lehmann, D., Ozaki, H., Pal, I., 1987. EEG alpha map series: brain micro-states by space-oriented adaptive segmentation. Electroencephalogr. Clin. Neurophysiol. 67, 271–288. https://doi.org/10.1016/0013-4694(87)90025-3

Little, S., Brown, P., 2014. The functional role of beta oscillations in Parkinson’s disease. Parkinsonism Relat. Disord. 20 Suppl 1, S44–48. https://doi.org/10.1016/S1353-8020(13)70013-0

Lopes, R., Delmaire, C., Defebvre, L., Moonen, A.J., Duits, A.A., Hofman, P., Leentjens, A.F.G., Dujardin, K., 2017. Cognitive phenotypes in parkinson’s disease differ in terms of brain-network organization and connectivity. Hum. Brain Mapp. 38, 1604–1621. https://doi.org/10.1002/hbm.23474

Lundqvist, M., Rose, J., Herman, P., Brincat, S.L., Buschman, T.J., Miller, E.K., 2016. Gamma and Beta Bursts Underlie Working Memory. Neuron 90, 152–164. https://doi.org/10.1016/j.neuron.2016.02.028

Merker, B.H., 2016. Cortical Gamma Oscillations: Details of Their Genesis Preclude a Role in Cognition. Front. Comput. Neurosci. 10, 78. https://doi.org/10.3389/fncom.2016.00078

Mheich, A., Hassan, M., Khalil, M., Gripon, V., Dufor, O., Wendling, F., 2018. SimiNet: A Novel Method for Quantifying Brain Network Similarity. IEEE Trans. Pattern Anal. Mach. Intell. 40, 2238–2249. https://doi.org/10.1109/TPAMI.2017.2750160

Michel, C.M., Koenig, T., 2018. EEG microstates as a tool for studying the temporal dynamics of whole-brain neuronal networks: A review. NeuroImage, Brain Connectivity Dynamics 180, 577–593. https://doi.org/10.1016/j.neuroimage.2017.11.062

Mørup, M., Hansen, L.K., 2009. Automatic relevance determination for multi-way models. J. Chemom. 23, 352–363. https://doi.org/10.1002/cem.1223

Nasreddine, Z.S., Phillips, N.A., Bédirian, V., Charbonneau, S., Whitehead, V., Collin, I., Cummings, J.L., Chertkow, H., 2005. The Montreal Cognitive Assessment, MoCA: A Brief Screening Tool For Mild Cognitive Impairment. J. Am. Geriatr. Soc. 53, 695–699. https://doi.org/10.1111/j.1532-5415.2005.53221.x

Nugent, A.C., Robinson, S.E., Coppola, R., Furey, M.L., Zarate, C.A., 2015. Group differences in MEG-ICA derived resting state networks: Application to major depressive disorder. NeuroImage 118, 1–12. https://doi.org/10.1016/j.neuroimage.2015.05.051

O’Neill, G.C., Tewarie, P., Vidaurre, D., Liuzzi, L., Woolrich, M.W., Brookes, M.J., 2018. Dynamics of large-scale electrophysiological networks: A technical review. NeuroImage, Brain Connectivity Dynamics 180, 559–576. https://doi.org/10.1016/j.neuroimage.2017.10.003

O’Neill, G.C., Tewarie, P.K., Colclough, G.L., Gascoyne, L.E., Hunt, B.A.E., Morris, P.G., Woolrich, M.W., Brookes, M.J., 2017. Measurement of dynamic task related functional networks using MEG. NeuroImage 146, 667–678. https://doi.org/10.1016/j.neuroimage.2016.08.061

Oswal, A., Brown, P., Litvak, V., 2013. Synchronized neural oscillations and the pathophysiology of Parkinson’s disease. Curr. Opin. Neurol. 26, 662–670. https://doi.org/10.1097/WCO.0000000000000034

Panagiotaropoulos, T., Kapoor, V., Logothetis, N., 2013. Desynchronization and rebound of beta oscillations during conscious and unconscious local neuronal processing in the macaque lateral prefrontal cortex. Front. Psychol. 4, 603. https://doi.org/10.3389/fpsyg.2013.00603

Pineda-Pardo, J.Á., Martínez, K., Solana, A.B., Hernández-Tamames, J.A., Colom, R., Pozo, F. del, 2015. Disparate Connectivity for Structural and Functional Networks is Revealed When Physical Location of the Connected Nodes is Considered. Brain Topogr. 28, 187–196. https://doi.org/10.1007/s10548-014-0393-3

Praamstra, P., Plat, E.M., Meyer, A.S., Horstink, M.W., 1999. Motor cortex activation in Parkinson’s disease: dissociation of electrocortical and peripheral measures of response generation. Mov. Disord. Off. J. Mov. Disord. Soc. 14, 790–799.

Praamstra, P., Stegeman, D.F., Cools, A.R., Horstink, M.W., 1998. Reliance on external cues for movement initiation in Parkinson’s disease. Evidence from movement-related potentials. Brain J. Neurol. 121 ( Pt 1), 167–177.

R Core Team, 2020. R: A Language and Environment for Statistical Computing. R Foundation for Statistical Computing, Vienna, Austria.

Ray, S., Maunsell, J.H.R., 2015. Do gamma oscillations play a role in cerebral cortex? Trends Cogn. Sci. 19, 78–85. https://doi.org/10.1016/j.tics.2014.12.002

Ridderinkhof, K.R., others, 2002. Activation and suppression in conflict tasks: Empirical clarification through distributional analyses.

Ridderinkhof, Richard K, K., Forstmann, B.U., Wylie, S.A., Burle, B., van den Wildenberg, W.P., 2011. Neurocognitive mechanisms of action control: resisting the call of the Sirens. Wiley Interdiscip. Rev. Cogn. Sci. 2, 174–192.

Rutledge, D.N., Jouan-Rimbaud Bouveresse, D., 2013. Independent Components Analysis with the JADE algorithm. TrAC Trends Anal. Chem. 50, 22–32. https://doi.org/10.1016/j.trac.2013.03.013

Sani, I., Stemmann, H., Caron, B., Bullock, D., Stemmler, T., Fahle, M., Pestilli, F., Freiwald, W.A., 2021. The human endogenous attentional control network includes a ventro-temporal cortical node. Nat. Commun. 12, 360. https://doi.org/10.1038/s41467-020-20583-5

Schmidt, R., Ruiz, M.H., Kilavik, B.E., Lundqvist, M., Starr, P.A., Aron, A.R., 2019. Beta Oscillations in Working Memory, Executive Control of Movement and Thought, and Sensorimotor Function. J. Neurosci. 39, 8231–8238. https://doi.org/10.1523/JNEUROSCI.1163-19.2019

Schmiedt-Fehr, C., Schwendemann, G., Herrmann, M., Basar-Eroglu, C., 2007. Parkinson’s disease and age-related alterations in brain oscillations during a Simon task. Neuroreport 18, 277–281. https://doi.org/10.1097/WNR.0b013e32801421e3

Simon, J.R., Rudell, A.P., 1967. Auditory SR compatibility: the effect of an irrelevant cue on information processing. J. Appl. Psychol. 51, 300.

Singer, W., 1999. Neuronal Synchrony: A Versatile Code for the Definition of Relations? Neuron 24, 49–65. https://doi.org/10.1016/S0896-6273(00)80821-1

Singh, A., Richardson, S.P., Narayanan, N., Cavanagh, J.F., 2018. Mid-frontal theta activity is diminished during cognitive control in Parkinson’s disease. Neuropsychologia 117, 113–122. https://doi.org/10.1016/j.neuropsychologia.2018.05.020

Sizemore, A.E., Bassett, D.S., 2018. Dynamic graph metrics: Tutorial, toolbox, and tale. NeuroImage, Brain Connectivity Dynamics 180, 417–427. https://doi.org/10.1016/j.neuroimage.2017.06.081

Skidmore, F., Korenkevych, D., Liu, Y., He, G., Bullmore, E., Pardalos, P.M., 2011. Connectivity brain networks based on wavelet correlation analysis in Parkinson fMRI data. Neurosci. Lett. 499, 47–51. https://doi.org/10.1016/j.neulet.2011.05.030

Smith, A., 1982. Symbol Digit Modalities Test (SDMT). Manual (revised). Los Angel. West. Psychol. Serv.

Spitzer, B., Haegens, S., 2017. Beyond the Status Quo: A Role for Beta Oscillations in Endogenous Content (Re)Activation. eNeuro 4, ENEURO.0170-17.2017. https://doi.org/10.1523/ENEURO.0170-17.2017

Stroop, J.R., 1935. Studies of interference in serial verbal reactions. J. Exp. Psychol. 18, 643.

Swann, N., Tandon, N., Canolty, R., Ellmore, T.M., McEvoy, L.K., Dreyer, S., DiSano, M., Aron, A.R., 2009. Intracranial EEG Reveals a Time- and Frequency-Specific Role for the Right Inferior Frontal Gyrus and Primary Motor Cortex in Stopping Initiated Responses. J. Neurosci. 29, 12675–12685. https://doi.org/10.1523/JNEUROSCI.3359-09.2009

Tabbal, J., Kabbara, A., Khalil, M., Benquet, P., Hassan, M., 2021. Dynamics of task-related electrophysiological networks: a benchmarking study. NeuroImage 231, 117829. https://doi.org/10.1016/j.neuroimage.2021.117829

Tadel, F., Baillet, S., Mosher, J.C., Pantazis, D., Leahy, R.M., 2011. Brainstorm: A User-Friendly Application for MEG/EEG Analysis. Comput. Intell. Neurosci. 2011, 879716. https://doi.org/10.1155/2011/879716

Tewarie, P., Liuzzi, L., O’Neill, G.C., Quinn, A.J., Griffa, A., Woolrich, M.W., Stam, C.J., Hillebrand, A., Brookes, M.J., 2019. Tracking dynamic brain networks using high temporal resolution MEG measures of functional connectivity. NeuroImage 200, 38–50. https://doi.org/10.1016/j.neuroimage.2019.06.006

Timmerman, M.E., Kiers, H.A.L., 2000. Three-mode principal components analysis: Choosing the numbers of components and sensitivity to local optima. Br. J. Math. Stat. Psychol. 53, 1–16. https://doi.org/10.1348/000711000159132

Tomlinson, C.L., Stowe, R., Patel, S., Rick, C., Gray, R., Clarke, C.E., 2010. Systematic review of levodopa dose equivalency reporting in Parkinson’s disease. Mov. Disord. Off. J. Mov. Disord. Soc. 25, 2649–2653. https://doi.org/10.1002/mds.23429

van den Wildenberg, W.P.M., Wylie, S.A., Forstmann, B.U., Burle, B., Hasbroucq, T., Ridderinkhof, K.R., 2010. To head or to heed? Beyond the surface of selective action inhibition: a review. Front. Hum. Neurosci. 4, 222. https://doi.org/10.3389/fnhum.2010.00222

van Wouwe, N.C., van den Wildenberg, W.P.M., Claassen, D.O., Kanoff, K., Bashore, T.R., Wylie, S.A., 2014. Speed pressure in conflict situations impedes inhibitory action control in Parkinson’s disease. Biol. Psychol. 101, 44–60. https://doi.org/10.1016/j.biopsycho.2014.07.002

Ville, D.V.D., Britz, J., Michel, C.M., 2010. EEG microstate sequences in healthy humans at rest reveal scale-free dynamics. Proc. Natl. Acad. Sci. 107, 18179–18184. https://doi.org/10.1073/pnas.1007841107

Wang, D., Zhu, Y., Ristaniemi, T., Cong, F., 2018. Extracting multi-mode ERP features using fifth-order nonnegative tensor decomposition. J. Neurosci. Methods 308, 240–247. https://doi.org/10.1016/j.jneumeth.2018.07.020

Wang, X.-J., 2010. Neurophysiological and computational principles of cortical rhythms in cognition. Physiol. Rev. 90, 1195–1268. https://doi.org/10.1152/physrev.00035.2008

Wechsler, D., 1981. WAIS-R Manual: Wechsler Adult Intelligence Scale-revised. Psychological Corporation.

Widge, A.S., Heilbronner, S.R., Hayden, B.Y., 2019. Prefrontal cortex and cognitive control: new insights from human electrophysiology. F1000Research 8, F1000 Faculty Rev-1696. https://doi.org/10.12688/f1000research.20044.1

Wiesman, A.I., Koshy, S.M., Heinrichs-Graham, E., Wilson, T.W., 2020. Beta and gamma oscillations index cognitive interference effects across a distributed motor network. NeuroImage 213, 116747. https://doi.org/10.1016/j.neuroimage.2020.116747

Wiesman, A.I., Wilson, T.W., 2020. Posterior Alpha and Gamma Oscillations Index Divergent and Superadditive Effects of Cognitive Interference. Cereb. Cortex 30, 1931–1945. https://doi.org/10.1093/cercor/bhz214

Winkler, A.M., Ridgway, G.R., Webster, M.A., Smith, S.M., Nichols, T.E., 2014. Permutation inference for the general linear model. NeuroImage 92, 381–397. https://doi.org/10.1016/j.neuroimage.2014.01.060

Wittfoth, M., Buck, D., Fahle, M., Herrmann, M., 2006. Comparison of two Simon tasks: Neuronal correlates of conflict resolution based on coherent motion perception. NeuroImage 32, 921–929. https://doi.org/10.1016/j.neuroimage.2006.03.034

Wolters, A.F., van de Weijer, S.C.F., Leentjens, A.F.G., Duits, A.A., Jacobs, H.I.L., Kuijf, M.L., 2019. Resting-state fMRI in Parkinson’s disease patients with cognitive impairment: A meta-analysis. Parkinsonism Relat. Disord. 62, 16–27. https://doi.org/10.1016/j.parkreldis.2018.12.016

Wu, H.-M., Hsiao, F.-J., Chen, R.-S., Shan, D.-E., Hsu, W.-Y., Chiang, M.-C., Lin, Y.-Y., 2019. Attenuated NoGo-related beta desynchronisation and synchronisation in Parkinson’s disease revealed by magnetoencephalographic recording. Sci. Rep. 9, 7235. https://doi.org/10.1038/s41598-019-43762-x

Wylie, S.A., Ridderinkhof, K.R., Bashore, T.R., van den Wildenberg, W.P.M., 2010. The effect of Parkinson’s disease on the dynamics of on-line and proactive cognitive control during action selection. J. Cogn. Neurosci. 22, 2058–2073. https://doi.org/10.1162/jocn.2009.21326

Wylie, S.A., Stout, J.C., Bashore, T.R., 2005. Activation of conflicting responses in Parkinson’s disease: evidence for degrading and facilitating effects on response time. Neuropsychologia 43, 1033–1043. https://doi.org/10.1016/j.neuropsychologia.2004.10.008

Wylie, S.A., van den Wildenberg, W.P.M., Ridderinkhof, K.R., Bashore, T.R., Powell, V.D., Manning, C.A., Wooten, G.F., 2009a. The effect of Parkinson’s disease on interference control during action selection. Neuropsychologia 47, 145–157.

Wylie, S.A., Van Den Wildenberg, W.P.M., Ridderinkhof, K.R., Bashore, T.R., Powell, V.D., Manning, C.A., Wooten, G.F., 2009b. The effect of speed-accuracy strategy on response interference control in Parkinson’s disease. Neuropsychologia 47, 1844–1853.

Yaesoubi, M., Miller, R.L., Calhoun, V.D., 2015. Mutually temporally independent connectivity patterns: A new framework to study the dynamics of brain connectivity at rest with application to explain group difference based on gender. NeuroImage 107, 85–94. https://doi.org/10.1016/j.neuroimage.2014.11.054

Yeung, N., Bogacz, R., Holroyd, C.B., Nieuwenhuis, S., Cohen, J.D., 2007. Theta phase resetting and the error-related negativity. Psychophysiology 44, 39–49. https://doi.org/10.1111/j.1469-8986.2006.00482.x

Zhu, Y., Liu, J., Ye, C., Mathiak, K., Astikainen, P., Ristaniemi, T., Cong, F., 2020. Discovering dynamic task-modulated functional networks with specific spectral modes using MEG. NeuroImage 218, 116924. https://doi.org/10.1016/j.neuroimage.2020.116924

